# MPK4 mediated phosphorylation of PIF4 controls thermosensing by regulation of H2A.Z deposition in Arabidopsis

**DOI:** 10.1101/2023.06.30.547269

**Authors:** Neetu Verma, Dhanraj Singh, Lavanya Mittal, Gopal Banerjee, Stanzin Noryang, Alok Krishna Sinha

## Abstract

Plants have the ability to perceive a slight upsurge in ambient temperature and respond by undergoing morphological changes, such as elongated hypocotyls and early flowering. The dynamic functioning of PHYTOCHROME INTERACTING FACTOR4 (PIF4) in thermomorphogenesis has been well established, although the regulatory pathway involved in thermosensing is not deciphered completely. In our study, we demonstrate that an increase in temperature from 22℃ to 28℃ induces the phosphorylation of PIF4 by MITOGEN-ACTIVATED PROTEIN KINASE 4 (MPK4) which shows high expression and activation at 28℃. Apparently, phosphorylated PIF4 represses the expression of *ACTIN-RELATED PROTEIN 6* (*ARP6*) that is required for mediating histone variant H2A.Z deposition at its target gene loci. We demonstrate that variation of *ARP6* expression in PIF4 phosphor -null and phosphor-mimetic seedlings affects hypocotyl growth and flowering at 22℃ and 28℃. Further, we show that change in *ARP6* expression affects H2A.Z deposition at the loci of genes involved in hypocotyl elongation using PIF4 phosphor -null and phosphor-mimetic seedlings. Interestingly, the expression of *MPK4* is also controlled by H2A.Z deposition in temperature dependent manner. Taken together, our findings highlight the cumulative molecular interplay between MPK4, PIF4, and chromatin modification by ARP6-mediated H2A.Z deposition as a regulatory mechanism of thermosensing.

## Introduction

Temperature is one of the most important determining factors in plant growth and development. Elevated temperature induces morphological changes such as elongated hypocotyl, hyponastic growth and early transition to flowering collectively known as thermomorphogenesis. These adaptive responses enhance the plant’s ability to survive and thrive under high temperature conditions. Thermomorphogenesis is instigated when plants perceive a change in temperature by thermosensing mechanism, a multifaceted process that is yet to be deciphered completely. The bHLH transcription factor PHYTOCHROME INTERACTING FACTOR 4 (PIF4) has been shown to play an important role in elevated temperature-induced thermomorphogenesis in *Arabidopsis thaliana* (Koini et al., 2009; Oh et al., 2012; Sun et al., 2012; Quint et al., 2016; Gangappa et al., 2017; Casal & Balasubramanian, 2019; Kim et al., 2020). PIF4 regulates the expression of many thermoresponsive genes such as *YUCCA8* (*YUC8*), *INDOLE-3-ACETIC ACID INDUCIBLE 19* (*IAA19*), *FLOWERING LOCUS T* (*FT*) and *PLANT U-BOX TYPE E3 UBIQUITIN LIGASE 31* (*PUB31*) (Frannklin et al., 2016; Sun et al., 2012; Kumar et al., 2016; Zhou et al., 2023). PIF4 activates the expression of *YUC8* and *IAA19*, while represses the expression *PUB31* in high temperature induced hypocotyl elongation (Frannklin et al., 2016; Sun et al., 2012; Kumar et al., 2016; Zhou et al., 2023). In addition to the PIF4 mediated transcriptional regulation, several reports show prominent role of epigenetic regulation in the thermosensing mechanism (Kumar & Wigge, 2010; Liu et al., 2015; Cortijo et al., 2017; Asensi-Fabado et al., 2017; Tian et al., 2019; Cui et al., 2021). The histone variant, H2A.Z exhibits temperature-dependent deposition around the transcription start site (TSS) of its target genes, leading to their transcriptional repression by wrapping the DNA more tightly around TSS (Mizuguchi et al., 2004; Kumar &Wigge, 2010; Cortijo et al., 2017). The role of ACTIN-RELATED PROTEIN 6 (ARP6) has been revealed in H2A.Z deposition-mediated temperature sensing in Arabidopsis (Kumar & Wigge, 2010; Cortijo et al., 2017). ARP6 is an important subunit of chromatin remodelling complex, SWR1 which is necessary for the insertion of H2A.Z into the nucleosomes of target gene loci at ambient temperature (Mizuguchi et al., 2004; March-Diaz and Reyes, 2009; Kumar & Wigge, 2010; Cortijo et al., 2017). Notably, *arp6* mutants exhibit increased expression of thermoresponsive genes like *HEAT SHOCK PROTEIN 70* (*HSP70*) and *FT* at ambient temperature due to the loss of H2A.Z occupancy at these gene loci (Kumar & Wigge, 2010; Gómez-Zambrano et al., 2019; Perrella et al., 2022).

PIF4 protein has been reported to undergo post-translational modifications in several signaling pathways. In red light signaling, phyB-induced phosphorylation leads to PIF4 degradation via the 26S proteasome pathway (Lorrain et al., 2008). Unlike red light, Blue light induced CRY1 interacts with PIF4 and suppresses its transcriptional activity (Ma et al., 2016; Pedmale et al., 2016). The role of post-translational modification of PIF4 has also been implicated in thermomorphogenesis (Foreman et al., 2011; Bernardo-Garcia et al., 2014; Choi et al., 2016; Han et al., 2022). Some recent reports suggest that SPA1 protein phosphorylates and stabilizes PIF4 at high temperature thereby regulating thermoresponsive growth (Lee et al., 2020; 2021). As post-translational modification of PIF4 is essential for its function in diverse pathways, therefore identification of other regulators of PIF4 is important to untangle the PIF4 mediated thermosensing mechanism entirely.

The role of MITOGEN-ACTIVATED PROTEIN KINASES (MAPKs) such as MPK3, MPK6 and MPK4 have also been implicated in plant responses to high temperature (Perez-Salamo et al., 2014; Andrasi et al., 2019). MAPK signaling cascade comprising of MAPK kinase kinases (MAPKKKs), MAPK kinases (MAPKKs) and MAPK, transduces the environmental signals to elicit intracellular responses (Sinha et al., 2011; Manna et al. 2023). MPK3/6 signaling promotes the production of guard cells in a response to high temperature (Lee et al., 2016). Tomato SlMPK1, a homologue of Arabidopsis MPK6, is reported to be a negative regulator of heat stress response (Ding et al., 2018). Recently a report suggests that MPK4 phosphorylates heat shock factor HSFA4A and modulates its activity leading to the induction of transcription of its target genes (Andrasi et al., 2019). However, the role of MAPK signaling in the PIF4 mediated thermosensing mechanism has not been elucidated so far.

Our study reveals an intriguing interplay of MPK4, PIF4 and ARP6/H2A.Z in thermosensing and hypocotyl growth in Arabidopsis. MPK4 interacts with and phosphorylates PIF4 at an elevated temperature of 28℃ and then phosphorylated PIF4 represses the expression of *ARP6* at 28℃. Moreover, our findings suggest that repression of *ARP6* at 28℃ leads to the eviction of H2A.Z from the loci of genes involved in hypocotyl elongation. Further, we demonstrate the role of ARP6 mediated H2A.Z insertion in the regulation of *MPK4* expression in a temperature dependent manner. Thus, our results establish that an interplay among PIF4, ARP6 and MPK4 regulates the temperature dependent plant traits such as hypocotyl growth and flowering.

## Results

### Establishing genetic relationship between PIF4 and ARP6

Firstly, we assessed the expression of *PIF4* and *ARP6* in *pif4* and *arp6* mutants to confirm their respective genotypes (Supplemental Fig. S1, A and B). To investigate the genetic relationship between PIF4 and ARP6, the WT, *pif4*, *arp6* and *PIF4*-OE plants were grown continuously either at ambient temperature of 22℃ or at elevated temperature of 28℃ under short-day (SD) conditions for 10 days. Interestingly, we observed that at ambient temperature of 22℃, *arp6* seedlings showed long hypocotyl phenotypes in comparison to that of WT, *pif4* and *PIF4*-OE seedlings (Fig. 1, A and B). In contrast, at the elevated temperature of 28℃, *pif4* mutant seedlings displayed shorter hypocotyl phenotypes compared to WT, *arp6*, and *PIF4*-OE seedlings (Fig. 1, C and D). This data indicated an antagonistic function of *PIF4* and *ARP6* in hypocotyl elongation. To further elaborate on this finding, the expression of *ARP6* and *PIF4* was studied in 10 days old WT seedlings grown at four different temperatures of 18℃, 22℃, 25℃ and 28℃ under short-day (SD) conditions. The expression of *ARP6* significantly decreased in seedlings grown at elevated temperatures of 25℃ and 28℃ (Supplemental Fig. S1C and Fig. 1E). Conversely, the expression of *PIF4* was significantly higher in seedlings grown at 25℃ and 28℃ compared to those grown at 18℃ and 22℃ (Supplemental Fig. S1C and Fig. 1E). Based on the antagonistic role of *PIF4* and *ARP6* in hypocotyl elongation and their expression patterns at different temperatures, we hypothesized that one might regulate other’s expression in a temperature-dependent manner. To investigate the potential regulatory relationship between *PIF4* and *ARP6*, we examined the transcript levels of *ARP6* and *PIF4* in *pif4* and *arp6* mutants, respectively, under both ambient and elevated temperature conditions. *PIF4* did not show any difference in the expression pattern between WT and *arp6* seedlings grown at different temperatures (Supplemental Fig. S1D). Therefore, it is unlikely that the expression of *PIF4* is regulated by ARP6/SWR1 mediated insertion of histone variant H2A.Z. When we examined the expression of *ARP6* in the *pif4* seedlings, we found that its expression increased at elevated temperatures of 25 and 28℃ compared to WT seedlings grown under the same conditions (Supplemental Fig. S1E and Fig. 1F). This observation suggests that *PIF4* may negatively regulate the expression of *ARP6*, particularly at elevated temperatures. To investigate whether high expression of *PIF4* at elevated temperatures is responsible for reduced expression of *ARP6*, we examined the expression of *ARP6* in *PIF4*-OE lines grown at different temperatures. First, the expression level of *PIF4* was analyzed in *PIF4*-OE seedlings grown at 18℃, 22℃, 25℃ and 28℃ (Supplemental Fig. S1F). *PIF4*-OE seedlings showed an increase in expression level of *PIF4* in comparison to WT seedlings. However, the expression of *ARP6* was significantly reduced in *PIF4*-OE seedlings at elevated temperatures of 25℃ and 28℃ (Supplemental Fig. S1G and Fig. 1F). Surprisingly, *PIF4*-OE lines did not show any decrease in the expression of *ARP6* at low and ambient temperatures of 18℃ and 22℃ when compared to WT (Supplemental Fig. S1G and Fig. 1F) suggesting the role of PIF4 in *ARP6* repression remarkably at elevated temperature. To further confirm that PIF4 is required for the suppression of *ARP6* promoter activity at elevated temperature, we generated transgenics in WT and *pif4* plants using *GUS* reporter gene under *ARP6* promoter. We observed a significant decline in GUS activity in case of WT-p*ARP6*:*GUS* transgenic seedlings grown at elevated temperature of 28℃ when compared to the seedlings grown at ambient temperature of 22℃ (Fig. 1, G and H). While in case of *pif4*-p*ARP6*:*GUS* transgenic seedlings, no decline in the GUS activity was observed at elevated temperature (Fig. 1, G and H). Therefore, all these data highlight that the expression of *ARP6* is suppressed by PIF4 specifically at elevated temperature establishing the fact that PIF4 and ARP6 function antagonistically in the same genetic pathway to regulate the hypocotyl length in response to the change in temperature.

**Fig. 1.**
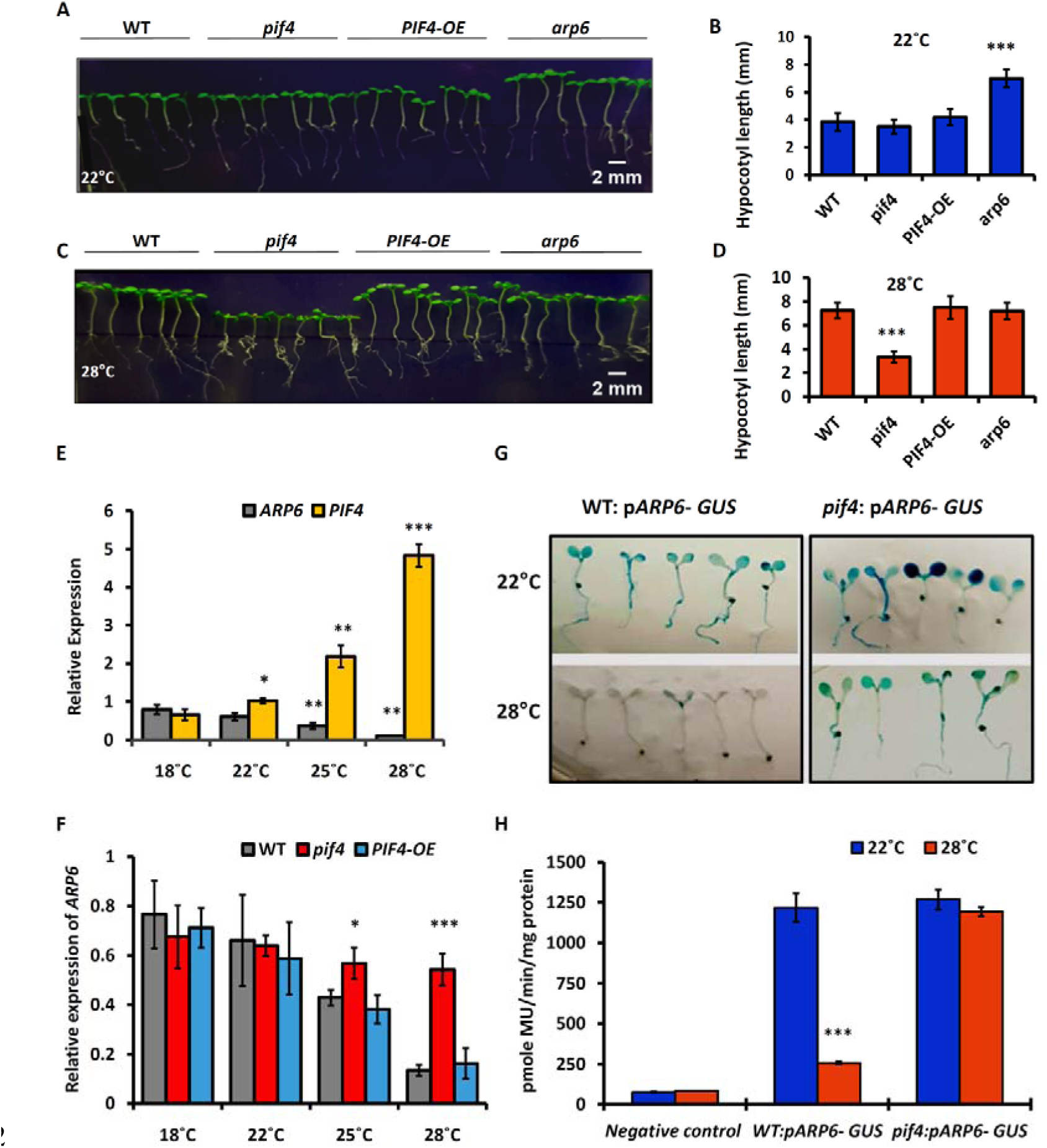
PIF4 promotes hypocotyl growth via repressing *ARP6* expression at elevated temperature. **(A)** Hypocotyl phenotypes of 10 days old seedlings grown under short days (SD) at 22℃. **(B)** Hypocotyl length of 10 days old seedlings grown at 22℃. Each bar represents mean ± s.d. (n = 20). Hypocotyl length data of each genotype (*pif4*, *PIF4*-OE and *arp6*) was compared to WT. One-Way ANOVA: ***P<0.0005 **(C)** Hypocotyl phenotypes of 10 days old seedlings grown under SD conditions at 28℃. **(D)** Hypocotyl length of 10 days old seedlings grown at 28 ℃. Each bar represents mean ± s.d. (n = 20). Hypocotyl length data of each genotype (*pif4*, *PIF4*-OE and *arp6*) was compared to WT. One-Way ANOVA: ***P<0.0005 **(E)** Relative transcript level of *ARP6* and *PIF4* in 10 days old WT seedlings at 4 temperatures. Each bar represents the mean ± s.d. (n= 3). The expression of *ARP6* and *PIF4* at 22℃, 25℃ and 28℃ was compared to their expression at 18℃. Student’s t test: *; *p* value < 0.05, **; *p* value < 0.005, ***; *p* value < 0.0005 **(F)** Relative transcript level of *ARP6* in 10 days old WT, *pif4* and *PIF4*-OE seedlings at 4 temperatures. Each bar represents mean ± s.d. (n= 3). *ARP6* expression in *pif4* and *PIF4*-OE seedlings was compared to WT. Student’s t test: *;*p* value < 0.05, ***; *p* value < 0.0005. **(G)** Histochemical GUS staining of 10 days old p*ARP6*-*GUS* transgenic seedlings grown at 22℃ and 28℃ under SD. WT:p*ARP6*-*GUS* and *pif4*:p*ARP6*-*GUS* represent the transgenic plants developed in WT and *pif4*, respectively. **(H)** Quantitative measurement of GUS activity in the WT:p*ARP6*-*GUS* and *pif4*:p*ARP6*-*GUS* seedlings at 22℃ and 28℃. Data at 28℃ was compared to 22℃. (n= 3) Student’s t test: (***; *P* value <0.0005).

### PIF4 represses the expression of *ARP6* by directly binding to its promoter

Further, we analyzed whether PIF4 regulates the expression of *ARP6* by directly binding to its promoter. Sequence analysis of p*ARP6-2* fragment identified two G-boxes (CANNTG) (Fig. 2A), the elements that are involved in binding to bHLH TFs (Oh et al., 2012; Zhang et al., 2013; Lau et al., 2018). Therefore, we initially performed *in-silico* docking to examine the interaction of PIF4 to *ARP6* promoter. To carry out the *in-silico* docking of PIF4 to the *ARP6* promoter, we required the 3D structures of the PIF4 protein and the *ARP6* promoter DNA. To obtain the structural information of the PIF4 protein, we used AlphaFold and predicted the overall structure of PIF4 (Supplemental Fig. S2A). A significant portion of the protein has an uncharacterized secondary structure with a low confidence score Supplemental Fig. S2, A and B. Therefore to ensure accurate docking of the PIF4 protein with the *ARP6* promoter, we specifically focused on predicting the structure of the DNA binding bHLH domain of PIF4 (Fig. 2, B and C). Predicted structure of PIF4 bHLH domain showed a high confidence score, as the residues are functionally conserved which is attributed to its DNA binding capability (Supplemental Fig. S2C). This approach allowed us to enhance the precision of our subsequent analysis and shed light on the potential interaction between PIF4 and the *ARP6* promoter. It is well known that PIF4, similar to many bHLH domain proteins, binds DNA as homo- and heterodimers (Littlewood and Evan, 1998; Toledo-Ortiz et al., 2003; Hornitschek et al., 2009; Hug and Qual, 2002; Gao et al., 2022). Therefore, we generated a dimer of the PIF4 bHLH domain using the PDB file of the structure of the bHLH domain predicted by Alpha fold and docking them with each other using HDock (Fig. 2D). The predicted structure of the PIF4 dimer was examined using PyMol to determine the amino acid residues responsible for the formation of the homodimer. Through careful analysis, we identified that the arginine residue R23 and the aspartic acid residue D44 are involved in the formation of homodimer (Fig. 2E). It is worth noting that the negatively charged side chains of aspartic acid form robust electrostatic bonds, commonly referred to as salt linkages, with the positively charged side group of arginine. Then, we performed docking of the PIF4 homodimer with the *ARP6* promoter using HDock. To accomplish this, we selected two 100bp *ARP6* promoter DNA fragments, each encompassing one of the G-boxes, for the docking with PIF4 homodimer. PIF4 homodimer exhibited binding to both G-boxes within the *ARP6* promoter with each monomer binding to complementary strands within the major groove of the DNA with a good confidence score (Fig. 2, F ^_^ I; Supplemental Fig. S2D). Further examination of docked structures using pymol revealed the key amino acid residues involved in this molecular interaction. Specifically, arginine (R14, R16), asparagine (N10), and histidine (H9) were identified as the crucial amino acid residues responsible for the molecular association of the PIF4 dimer with G-box motif 1 (CATTTG) within the *ARP6* promoter on both DNA strands (Fig. 2G). Furthermore, same amino acid residues (R14, R16, N10) were involved in the interaction of the homodimer with G-box motif 2 (CACCTG) (Fig. 2I). The only difference observed in this case was, instead of histidine (H9), the lysine residue (K39) participated in bond formation with a distinct nucleotide within the DNA (Fig. 2I). To further validate the results obtained from *in-silico* docking, we conducted a yeast one-hybrid assay to test whether PIF4 binds to the *ARP6* promoter *in vitro* using the three different fragments, p*ARP6*-1, p*ARP6*-2 & p*ARP6*-3 (each of ∼350 bp) of *ARP6* promoter (Fig. 2J). Yeast one-hybrid assays showed that PIF4 specifically interacted with p*ARP6*-2 fragment, while no interaction of PIF4 was observed with p*ARP6*-1 and p*ARP6*-3 fragments (Fig. 2J). Notably, both of the identified G-boxes are present within the *pARP6-2* fragment, which is in line with our *in-silico* analysis findings. Next, we performed electrophoretic mobility shift assays (EMSA) to further examine PIF4 binding to *ARP6* promoter *in vitro* using the DNA probes 1 and 2 containing G-box elements 1 and 2, respectively. Sequences of the WT and mutated probes have been depicted in Supplemental Fig. S2E. As expected, PIF4 showed interaction with both the probes, 1 and 2 (Fig. 2K). Mutation in G-box elements completely abolished the interaction of PIF4 to both probes (Fig. 2K). To confirm the PIF4 biding to the p*ARP6-2* fragment *in vivo*, we performed chromatin immunoprecipitation (ChIP) qPCR assay using seedlings of *PIF4* complementation line (*PIF4*-3X*HA*) which were generated by transforming the *pif4* mutant with *PIF4* fused with 3X*HA* tag driven by its native promoter. Equal number of *PIF4*-3X*HA* seedlings were grown for 10 days at ambient temperature of 22℃ and elevated temperature of 28℃, respectively. The immunoprecipitation of PIF4 protein using anti-hemagglutinin (HA) antibody revealed a higher abundance of PIF4 protein in *PIF4*-3X*HA* seedlings grown at 28℃ in comparison to seedlings grown at 22℃ (Fig. 2L). qPCR data showed that p*ARP6-1,* p*ARP6-2* and p*ARP6-3* fragments of *ARP6* promoter were significantly enriched in the immunoprecipitated fraction of *PIF4*-3X*HA* seedlings grown at elevated temperature of 28℃ in comparison to seedlings grown at 22℃ (Fig. 2M). Notably, the p*ARP6-2* fragment exhibited the highest level of enrichment among the three fragments in *PIF4*-3X*HA* seedlings. Together, these data demonstrate that PIF4 TF directly binds to the *ARP6* promoter.

**Fig. 2.**
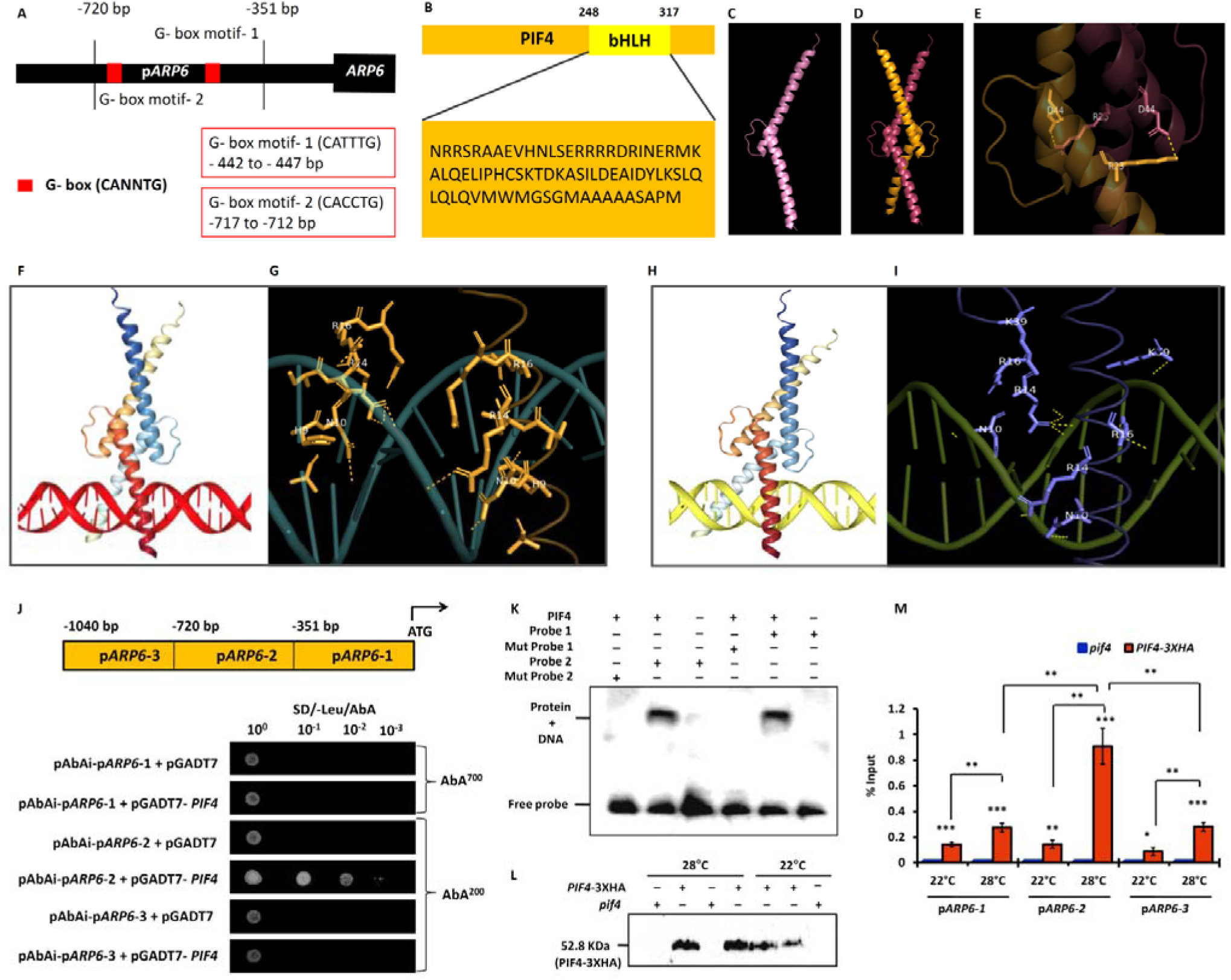
Direct binding of PIF4 TF to G boxes in *ARP6* promoter. **(A)** Illustration of G-box motifs in *ARP6* promoter. **(B)** Amino acid sequence of bHLH domain of PIF4 protein. **(C)** Predicted structure of bHLH domain of PIF4. **(D)** Predicted structure of the PIF4 bHLH homodimer. **(E)** Amino acid residues involved in the formation of PIF4 homodimer. **(F)** Predicted binding of PIF4 bHLH homodimer to G-box 1 of *ARP6* promoter. **(G)** Amino acid residues involved in interaction of PIF4 bHLH domain with the G-box 1. **(H)** Predicted binding of PIF4 bHLH homodimer to G-box 2 of *ARP6* promoter. **(I)** Amino acid residues involved in interaction of PIF4 bHLH domain with the G-box 2. **(J)** Upper panel is the **s**chematic of *ARP6* promoter showing the positions of p*ARP6*-1, p*ARP6*-2 & p*ARP6*-3 fragments used in yeast one-hybrid assay. Lower panel is the yeast one-hybrid assay showing the interaction of PIF4 TF with p*ARP6* fragments. **(K)** EMSA showing the interaction of PIF4 with *ARP6* promoter probes 1 and 2 containing G-box, 1 and 2, respectively. The experiment was repeated two times with similar results. **(L** and **M)** ChIP-qPCR assays performed in 10 days old *pif4* and *PIF4-3XHA* seedlings grown at 22 and 28℃ under SD using anti-HA antibody. (L) Immunoblotting of PIF4 in the immunoprecipitated chromatin using anti-HA antibody. (M) Analysis of interaction of p*ARP6-1*, p*ARP6-2* and p*ARP6-3* fragments with PIF4 TF by qPCR. Each bar represents mean ± s.d. of three technical replicates. Enrichment of p*ARP6* fragments at 28℃ was compared to 22℃. Enrichment of p*ARP6-2* was also compared to p*ARP6-1* and p*ARP6-3* at 28℃. Student’s t test. (*; *P* value <0.05, **; *P* value <0.005, ***; *p* value < 0.0005). Two independent experiments were performed with similar results.

### MPK4 upregulation, activation and interaction with PIF4 at elevated temperature

PIF4 is regulated by numerous post-translational modifications which affect its transcriptional activity and stability (Foreman et al., 2011; Bernardo-Garcia et al., 2014; Han et al., 2022). Given that *PIF4* specifically influences *ARP6* expression at elevated temperatures, we speculated that PIF4 may undergo temperature-dependent post-translational modification. MAP kinase, MPK4 was shown to play a regulatory role in Arabidopsis under heat stress (Andrasi et al., 2019). Therefore, we analyzed the expression of all 20 members of MAPK genes (*MPK1*-*MPK20*) in 10 days old Arabidopsis seedlings grown at ambient and elevated temperatures (Supplemental Fig. S3A). Three MAP kinase genes viz *MPK4*, *MPK7* and *MPK13* showed upregulation in the seedlings grown at 28℃. Among these three, the upregulation of *MPK4* was found to be more significant in comparison to *MPK7* and *MPK13*. Further building upon the lead, we examined the protein level of MPK4 via immunoblotting in the 10 days old WT, *mpk3*, *mpk4* and *mpk6* seedlings grown at 22℃ and 28℃ (Fig. 3A). We observed an increase in protein level of MPK4 in the WT, *mpk3* and *mpk6* seedlings grown at 28℃. To further investigate this observation, we generated transgenic plants using *GUS* reporter gene under *MPK4* promoter. 10 days old p*MPK4*-*GUS* transgenic seedlings grown at elevated temperature of 28℃ displayed higher GUS activity in comparison to seedlings grown at 22℃ (Fig. 3B). These data implicate that *MPK4* expression upregulates in response to elevated temperature. Next, we examined the MAPK activities in WT, *mpk3*, *mpk4* and *mpk6* seedlings grown at ambient and elevated temperatures using an anti-pTEpY antibody. Interestingly we observed an activation of MPK4 in response to elevated temperature in WT, *mpk3* and *mpk6* seedlings (Fig. 3C). Subsequently, we evaluated the role of MPK4 in hypocotyl growth at elevated temperature by observing the hypocotyl phenotype and length of *mpk4* mutant seedlings. *mpk4* seedlings grown at 28℃ showed shorter hypocotyl length in comparison to WT seedlings (Supplemental Fig. S3, B and C). *mpk4* seedlings showed hypocotyl phenotype similar to that of *pif4* seedlings at elevated temperature suggesting that PIF4 and MPK4 act in the same genetic pathway to control temperature dependent hypocotyl growth. All these data led us to examine the interaction between MPK4 and PIF4 proteins. Firstly the interaction between these two proteins was examined *in vitro* by Yeast two-hybrid assay. PIF4 showed strong interaction with MPK4 in both the instances when *MPK4* and *PIF4* were cloned into the pGBKT7 and pGADT7 vectors, respectively and also when these genes were swapped between the above vectors (Supplemental Fig. S3, D and E). However, no interaction of PIF4 was observed with MPK7 which is a close relative of MPK4 thus, indicating the specificity of PIF4-MPK4 interaction (Supplemental Fig. S3, D and E). Next, we performed the co-immunoprecipitation (co-IP) assay to analyze the interaction between these two proteins *in vivo* using *PIF4*-3X*HA* transgenic seedlings at 22℃ and 28℃. PIF4 protein was immunoprecipitated from *PIF4*-3X*HA* seedlings grown at two different temperatures of 22℃ and 28℃ using anti-HA antibody and the interaction of MPK4 in the immunoprecipitated PIF4 protein was analyzed by immunoblotting with anti-MPK4 antibody. A band of expected size of MPK4 was detected in the anti-HA immunoprecipitate of *PIF4*-3X*HA* seedlings grown at 28℃ (Fig. 3D). No MPK4 protein band was observed in the anti-HA immunoprecipitate of *PIF4*-3X*HA* seedlings grown at an ambient temperature of 22℃ (Fig. 3D). Another co-IP experiment was performed by immunoprecipitating MPK4 protein using anti-MPK4 antibody and interaction of PIF4 protein in the precipitated MPK4 protein fraction was monitored by immunoblotting with anti-HA antibody. PIF4 protein band was detected in the MPK4 fraction immunoprecipitated from *PIF4*-3X*HA* seedlings at 28℃ (Fig. 3E). From the result of co-IP expreriment, we can assume that after activation of MAP kinase activity at elevated temperature, MPK4 interacts with PIF4. To test this assumption we conducted a Bimolecular Fluorescence Complementation (BiFC) experiment in tobacco leaves. The leaves of *Nicotiana benthamiana*were infiltrated with the MPK4 and PIF4 constructs and the plants were grown at both 22 and 28℃ for 2 days before harvesting the leaves. We examined MPK4 activation in the infiltrated leaves at 22 and 28℃ using anti-pTpEY antibody. MPK4 protein was immunoprecipitated from the total protein extract of leaf halves. MPK4 activation was analyzed in the immunoprecipitated MPK4 protein using anti-pTpEY antibody. We observed MPK4 activation in the leaves taken from the plants grown at 28℃ (Fig. 3F). Then, we used the remaining leaf halves for monitoring interaction between MPK4 and PIF4 proteins at 22 and 28℃. We observed the interaction between these two proteins specifically at 28℃ (Fig. 3G). Altogether, these results provide compelling evidences that MPK4 is upregulated, activated and subsequently interacts with PIF4 at an elevated temperature of 28℃.

**Fig. 3.**
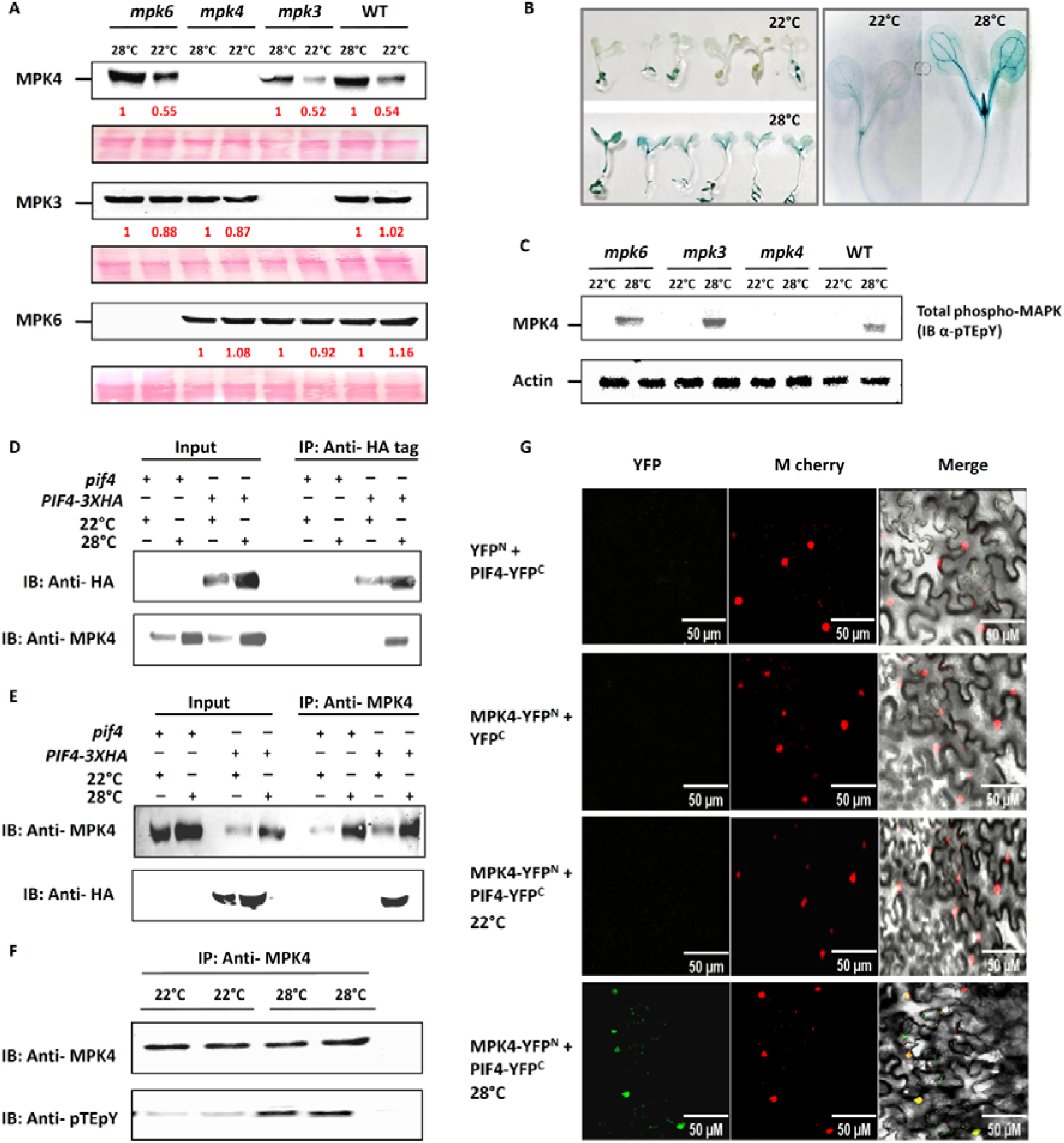
*MPK4* upregulation, activation and interaction with PIF4 at elevated temperature. **(A)** Immunoblotting of MPK4, MPK3 and MPK6 in 10 days old WT, *mpk3*, *mpk4* and *mpk6* seedlings grown at 22 and 28℃ under SD. Numbers in red indicate the intensity of protein bands. Band intensity at 22℃ is relative to band intensity at 28℃. Ponceau staining of rubisco was used as loading control. The experiment was repeated three times with similar results. **(B)** GUS staining of 10 days old p*MPK4*-*GUS* transgenic seedlings grown at 22℃ and 28℃. **(C)** Analysis of MAP Kinase activity in WT, *mpk4*, *mpk3* and *mpk6* seedlings at 22 and 28℃ using anti-pTEpY antibody. Immunoblotting of Actin was used as control **(D** and **E)** Co-IP assay showing interaction between PIF4 and MPK4 in temperature-dependent manner. (D) *PIF4-*3X*HA* seedlings grown at 22 and 28℃ were used for immunoprecipitation of PIF4-3XHA protein using anti-HA antibody. Total (left side) and precipitated (right side) proteins were analyzed by immunoblotting using antibodies against HA tag and MPK4, respectively. (E) *PIF4-*3X*HA* seedlings grown at 22 and 28℃ were used for immunoprecipitation of MPK4 protein using anti-MPK4 antibody. Total (left side) and precipitated (right side) proteins were analyzed by immunoblotting using anti-MPK4 and anti-HA antibodies, respectively. **(F and G)** BiFC experiment showing protein– protein interaction between PIF4 and MPK4 in *N. benthamiana*. (F) Analysis of MAP Kinase activity in the leaf tissue infiltrated with PIF4 and MPK4 constructs at 22 and 28℃. Protein extracts from the infiltrated leaves were used to immunoprecipitate MPK4 protein using anti-MPK4 antibody. Precipitated proteins were analyzed by immunoblotting using anti-MPK4 (upper panel) and anti-pTEpY (lower panel) antibodies. (G) BiFC experiment showing interaction between MPK4 and PIF4 specifically at 28℃ in the nucleus of leaf epidermal cells with YFP signal. Scale bar, 50µm.

### MPK4 phosphorylates PIF4 at elevated temperature

To investigate the potential phosphorylation of PIF4 by MPK4, we conducted an *in vitro* phosphorylation assay using His-tagged MPK4 and GST-tagged PIF4 proteins expressed in bacteria. We observed a strong phosphorylation of PIF4 by MPK4 as demonstrated in Fig. 4A. To confirm the phosphorylation of PIF4 *in vivo*, we performed a Phos-tag mobility shift assay. For the experiment, *PIF4*-3X*HA* seedlings were grown at 22℃ and 28℃ temperatures and protein extracts from these seedlings were separated in a phos-tag gel. Immunoblot analysis of PIF4 protein was then performed using anti-HA antibody. Remarkably, the PIF4 bands were shifted to higher molecular weight In the seedlings grown at elevated temperature (Fig. 4B). The role of MPK4 in PIF4 phosphorylation *in vivo* was further investigated by transient expression of *PIF4*-3X*HA* construct into the WT and *mpk4* seedlings. The seedlings were suspended into a medium containing *Agrobacterium* suspension harbouring the *PIF4*-3X*HA* construct and incubated at 22℃ for 36 hrs. After incubation, the seedlings were surface sterilised and grown for three more days at 22 and 28℃. The phosphorylation of PIF4 was monitored in the harvested seedlings by Phos-tag mobility shift assay. One band of up-shifted PIF4 protein was completely abolished in the *mpk4* seedlings at 28℃ in comparison to WT seedlings at 28℃ (Fig. 4B). The upshift of other PIF4 protein band was not completely abolished in *mpk4* seedlings at 28℃, suggesting that MPK4 is not the only kinase which is responsible for the phosphorylation of PIF4 protein at elevated temperature. These results indicate that MPK4 is one of the kinases that phosphorylates PIF4 at elevated temperature.

**Fig. 4.**
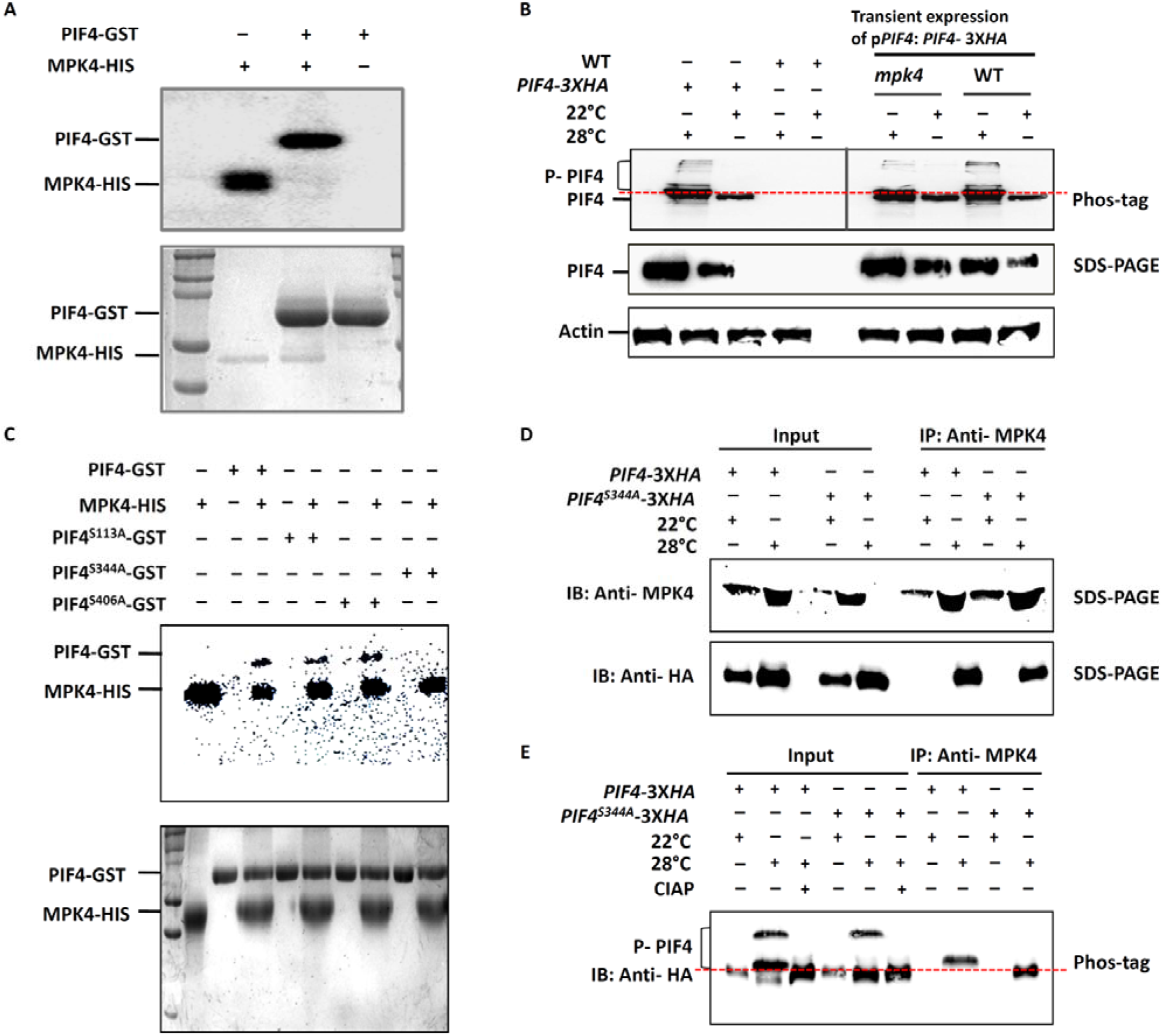
MPK4 phosphorylates PIF4 at elevated temperature. **(A)** *In vitro* phosphorylation assay using bacterially expressed PIF4-GST and MPK4-His recombinant proteins. PIF4 phosphorylation was detected by autoradiography after gel electrophoresis (top panel). Lower panel shows the Coomassie brilliant blue (CBB)-stained gel with the positions of different proteins indicated. **(B)** PIF4 phosphorylation at elevated temperature under *in vivo* condition. Left panel shows the up-shifted bands of phosphorylated PIF4 protein in *PIF4*-3X*HA* seedlings at 28℃ using anti-HA antibody. Right panel shows the phosphorylation of PIF4 in *mpk4* and WT seedlings transiently expressing *PIF4*-3X*HA* under the *PIF4* promoter at 22 & 28℃. Immunoblotting of Actin protein was used as a loading control. **(C)** *In-vitro* phosphorylation assay with three mutated PIF4 proteins, PIF4^S113A^, PIF4^S344A^ and PIF4^S406A^. The upper panel is the autoradiograph to analyze the phosphorylation status of mutated PIF4 proteins and the lower panel is the CBB stained gel with the positions of different proteins indicated. **(D and E)** Co-IP assay showing the interaction between phosphor-null PIF4^S344A^ and MPK4, and phosphorylation status of PIF4^S344A^ in a temperature-dependent manner. *PIF4-*3X*HA* and *PIF4*^S344A^-3X*HA* transgenic seedlings grown at 22 and 28℃ were used for immunoprecipitation of MPK4 protein using anti-MPK4 antibodies. Total (left side) and precipitated (right side) proteins were analyzed by (D) immunoblotting using anti-MPK4 and anti-HA antibodies, respectively; (E) immunoblotting using anti-HA antibodies to show the phosphorylation status of PIF4 and PIF4^S344A^ proteins on a phos-tag gel.

Next, we decided to identify the specific amino acid residue of PIF4 which is phosphorylated by MPK4. MAP kinases are known to phosphorylate their substrates at specific serine/threonine residues which are followed by a characteristic proline amino acid (Singh and Sinha, 2015). Based on this knowledge, we screened the protein sequence of PIF4 for the presence of serine/threonine residues followed by a proline amino acid. Analysis of PIF4 protein sequence showed three putative serine residues at 113^th^, 344^th^ and 406^th^ positions followed by a characteristic proline amino acid (Supplemental Fig. S4A). Additionally, we performed IP-MS assays using *PIF4*-3X*HA* seedlings grown at 22 and 28℃, respectively. *PIF4*-3X*HA* seedlings grown at 28℃, specifically showed phospho-peptide containing ser344 residue in the analysis (Supplemental Fig. S4B). Subsequently, we mutated 113^th^, 344^th^ and 406^th^ serine residues to alanine, a non-phosphorytable amino acid to generate three mutated versions of PIF4 protein named as PIF4^S113A^, PIF4^S344A^ and PIF4^S406A^. We performed the *in-vitro* phosphorylation assay using these mutated PIF4 proteins. PIF4^S113A^ and PIF4^S406A^ showed phosphorylation signal similar to that of WT PIF4 (Fig. 4C). Whereas, the phosphorylation of PIF4^S344A^ was completely abolished as shown in Fig. 4C suggesting that MPK4 phosphorylates PIF4 specifically at the serine-344 residue. To assess the role of serine 344 in the phosphorylation of PIF4 *in vivo*, we generated phosphor-null transgenic plants by transforming *pif4* plants with *PIF4*^S344A^ gene fused with HA tag under *PIF4* promoter. First, we immunoprecipitated PIF4 and PIF4^S344A^ proteins from *PIF4*-3X*HA* and *PIF4*^S344A^-3X*HA* seedlings, respectively at 22℃ and 28℃ using anti-HA antibody. Then, the immunoprecipitated PIF4 and PIF4^S344A^ proteins were analyzed by immunoblotting using anti-phospho-serine (anti-pSer) antibody. PIF4 protein band was observed in the immunoprecipitated PIF4 protein fraction at 28℃ by immunoblotting with anti-pSer antibody (Supplemental Fig. S4B). A very faint band of PIF4 ^S344A^ protein was observed in the immunoprecipitated PIF4^S344A^ protein fraction by immunoblotting with anti-pSer antibody (Supplemental Fig. S4B). This result shows the specificity of serine 344 residue in the phosphorylation of PIF4. Next, we performed a co-IP experiment to analyze the interaction between PIF4^S344A^ and MPK4 proteins and the role of serine 344 in the phosphorylation of PIF4 using *PIF4*-3X*HA* and *PIF4*^S344A^-3X*HA* transgenic seedlings grown at 22℃ and 28℃. MPK4 protein from *PIF4*-3X*HA*, *PIF4*^S344A^-3X*HA* seedlings was immunoprecipitated using anti-MPK4 antibody and interaction of PIF4^S344A^ protein in the precipitated MPK4 protein fraction was subsequently examined by immunoblotting with anti-HA antibody. PIF4 ^S344A^ protein band was detected in the MPK4 fraction immunoprecipitated from *PIF4*^S344A^-3X*HA* seedlings at 28℃ (Fig. 4D). This result indicates that replacement of serine 344 with alanine does not affect the interaction of PIF4 protein with MPK4. Another co-immunoprecipitations of PIF4 and PIF4^S344A^ proteins were performed using SDS phos-tag gel. Input and the co-immunoprecipitated fractions of *PIF4*-3X*HA* and PIF4^S344A^-3X*HA* seedlings at 22℃ and 28℃ were resolved over SDS phos-tag gel. Immunoprecipitation of input fraction of *PIF4*-3X*HA* using anti-HA antibody showed two bands of upshifted PIF4 proteins at 28℃, while in case of *PIF4*^S344A^-3X*HA* seedlings, only upper band of upshifted PIF4^S344A^ protein was visible, whereas the lower upshifted band was completely abolished at 28℃ (Fig. 4E) as observed in case of Fig. 4B. In case of co-immunoprecipitation of PIF4 and PIF4^S344A^ proteins from immunoprecipitated MPK4 fractions of *PIF4*-3X*HA* and *PIF4*^S344A^-3X*HA* seedlings, respectively, WT PIF4 protein showed an upshift in comparison to that of PIF4^S344A^ protein at 28℃ (Fig. 4E). Altogether, these data indicate that serine 344 is specifically involved in the phosphorylation of PIF4 by MPK4 at an elevated temperature of 28℃.

### MPK4-mediated phosphorylation of PIF4 is essential for the *ARP6* repression and induction of hypocotyl elongation and early flowering at elevated temperature

First of all, To determine that both MPK4 and PIF4 proteins are required for temperature dependent regulation of *ARP6* expression, we generated *pif4*/*mpk4* double mutant. The generation of *pif4*/*mpk4* double mutant has been described in detail in the materials and methods section. Firstly, we examined the hypocotyl phenotype and length of *pif4*/*mpk4* seedlings at 22 and 28℃ and compared it with WT, *pif4* and *mpk4* seedlings. At both temperatures, *pif4*/*mpk4* seedlings showed a drastically dwarf hypocotyl in comparison to WT, *pif4* and *mpk4* seedlings (Supplemental Fig. S5, A ^_^ D). Further, we monitored the expression of *ARP6* in *pif4*/*mpk4* double mutant seedlings at 22 and 28℃ and compared it with *ARP6* expression in *pif4*, *mpk4* and WT (Supplemental Fig. S5E). In comparison to WT seedlings, *pif4*/*mpk4* seedlings showed an increase in *ARP6* expression level at 28℃ similar to that of *pif4* and *mpk4* seedlings. These results suggest that *ARP6* regulation by the coordinated actions of MPK4 and PIF4 controls the thermoresponsive hypocotyl growth in Arabidopsis. Next, we performed transient expression assays into *pif4*, *mpk4* and *pif4*/*mpk4* seedlings using effector constructs (*PIF4* and *MPK4* under their native promoters) to determine the necessity of both MPK4 and PIF4 proteins for *ARP6* repression at 28℃. The schematic presentations of the effector constructs are shown in Fig. 5A. 7 days old 100 seedlings of each genotype were incubated with *Agrobacterium* cells carrying the effector constructs at 22℃ for 36 hrs. After infection, the seedlings were surface sterilized and washed with water. Subsequently, the seedlings were grown at 22 and 28℃ for 3 days before harvesting. Transient expression of PIF4 and MPK4 proteins were confirmed by immunoblotting of these proteins in 50 infected seedlings of each genotype with anti-HA and anti-MPK4 antibodies (Fig. 5, B ^_^ D). Then the expression of *ARP6* was examined in the rest of 50 infected *pif4*, *mpk4* and *pif4*/*mpk4* seedlings (Fig. 5, E ^_^ G). *ARP6* showed a reduction in transcript level at 28℃ in the *pif4* seedlings infected with the *PIF4* effector construct when compared to *pif4* seedlings infected with the empty vector construct (Fig. 5E). As well, *mpk4* seedlings infected with the *MPK4* effector construct at 28℃, showed reduction in *ARP6* transcript level when compared to *mpk4* seedlings infected with the empty vector construct (Fig. 5F). On the contrary, no transcript repression of *ARP6* was observed in *pif4*/*mpk4* seedlings infected with either *PIF4* or *MPK4* effector construct at 28℃ (Fig. 5, E and F). Repression of *ARP6* transcript was observed in the *pif4*/*mpk4* seedlings infected with both the *PIF4* and *MPK4* effector constructs at 28℃ (Fig. 5G). These data suggest that both MPK4 and PIF4 are required for the repression of *ARP6* gene expression at elevated temperature. Further, we explored the role of MPK4-mediated PIF4 phosphorylation in *ARP6* repression at elevated temperature. We monitored the *ARP6* expression in the WT *PIF4*-3X*HA,* phosphor-null *PIF4*^S344A^-3X*HA* and phosphor-mimetic *PIF4*^S344D^-3X*HA* seedlings at 22 and 28℃ (Fig. 5H). When compared to *PIF4*-3X*HA* seedlings, *PIF4*^S344A^-3X*HA* seedlings showed higher expression of *ARP6* at 28℃ similar to that of *pif4* seedlings (Fig. 5H). *PIF4*^S344D^-3X*HA* seedlings showed lower expression of *ARP6* in comparison to *PIF4*-3X*HA* seedlings at 22℃ (Fig. 5H). As we have earlier shown that MPK4 was unable to phosphorylate PIF4^S344A^ (Fig. 4, C and E), Therefore, the observed increase in *ARP6* expression in *PIF4*^S344A^-3X*HA* seedlings at 28℃ provided direct evidence of the necessity of PIF4 phosphorylation by MPK4 at elevated temperature for *ARP6* repression. Next, we were interested to determine whether mutation of serine 344 amino acid residue to alanine affects the binding of PIF4^S344A^ to the *ARP6* promoter. To investigate the possibility, we performed EMSA using bacterially expressed WT PIF4 and phosphor-null PIF4^S344A^ proteins with p*ARP6* probes. First, PIF4 and PIF4^S344A^ proteins were phosphorylated by incubating with MPK4 following which the phosphorylated proteins were incubated with p*ARP6* probes. Interestingly, we detected an upshift with both the phosphorylated and nonphosphorylated WT PIF4 protein, respectively (Supplemental Fig. S6). Additionaly, phosphor-null PIF4^S344A^ protein also showed upshift similar to that of WT PIF4 protein (Supplemental Fig. S6). To further substantiate this finding, we performed the ChIP-qPCR assays using *PIF4*-3X*HA* and*PIF4*^S344A^-3X*HA* seedlings grown at 22 and 28℃. The ChIP-qPCR assay demonstrated no significant difference in the association of PIF4^S344A^ protein to the p*ARP6*-2 fragment when compared to PIF4 protein at 22 and 28℃ (Fig. 5I), indicating that phosphorylation of PIF4 is not required for its binding to the *ARP6* promoter and represses the transcription of *ARP6* by a distinct mechanism.

**Fig. 5.**
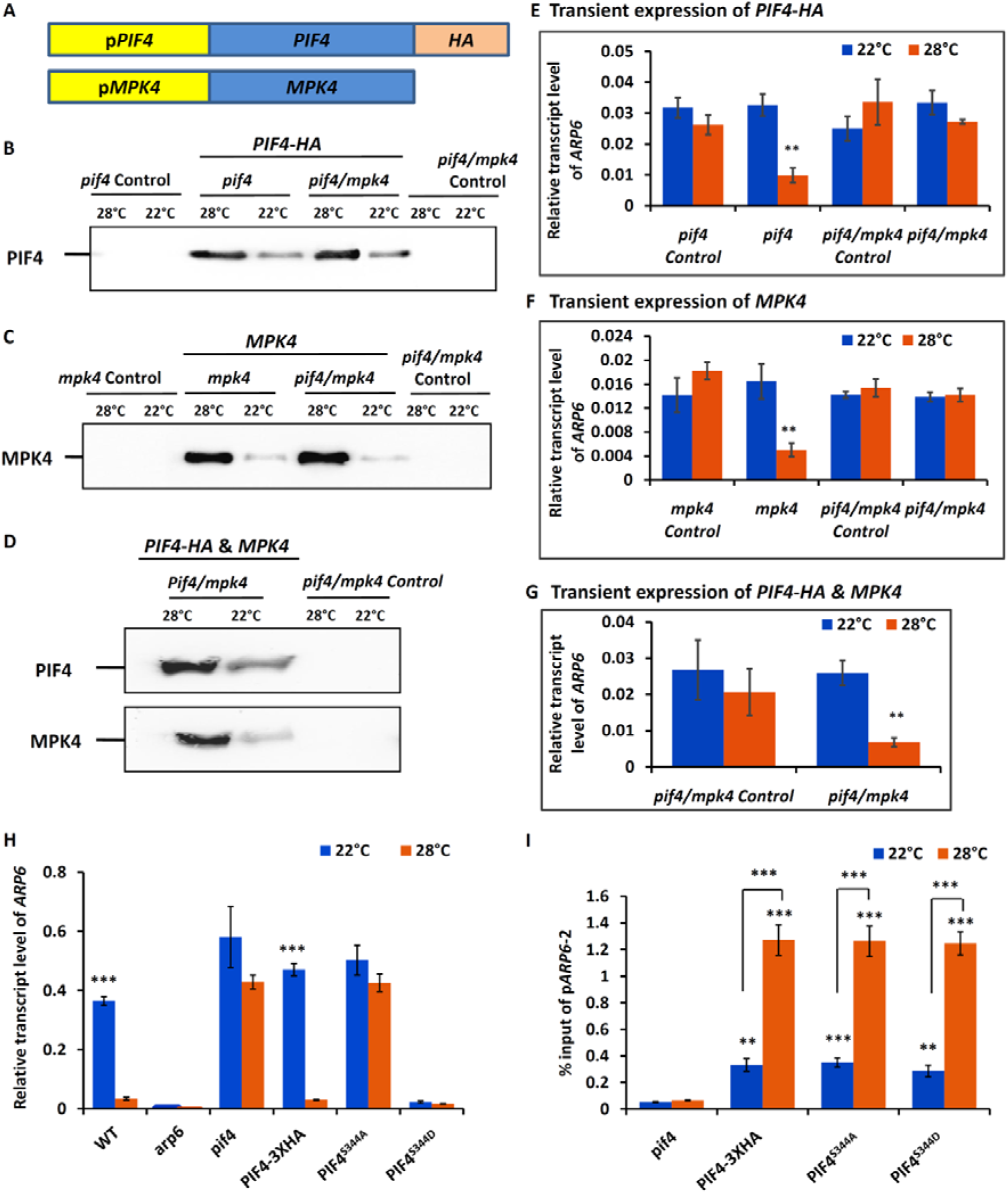
MPK4 mediated phosphorylation of *PIF4* is necessary for *ARP6* repression at elevated temperature. **(A)** Schematics of effector constructs used for the transient expression of *PIF4* and *MPK4* genes in Arabidopsis seedlings. **(B, C and D)** Confirmation of transient expression of *PIF4* in *pif4* and *pif4*/*mpk4* (B), *MPK4* in *mpk4* and *pif4*/*mpk4* (C) and *PIF4*, *MPK4* together in *pif4/mpk4* seedlings (D) by immunoblotting using anti-HA & anti-MPK4 antibodies. Control in each transient expression experiment denotes the infection of seedlings with *Agrobacterium* cells carrying an empty vector. **(E, F and G)** Quantification of *ARP6* transcript level in *pif4* and *pif4*/*mpk4* seedlings infected with *PIF4*-3X*HA* effector construct (E), *mpk4* and *pif4*/*mpk4* seedlings infected with *MPK4* effector construc t (F) and *pif4*/*mpk4* seedlings infected with both the *PIF4*-3X*HA* and *MPK4* effector constructs (G). Each bar represents the mean ± s.d. of three replicates. The expression of *ARP6* at 28℃ was compared to its expression at 22℃. Student’s t test. (**; *p* value < 0.005) **(H)** Transcript level of *ARP6* in 10 days old WT, *arp6*, *pif4*, *PIF4-*3X*HA*, *PIF4*^S344A^ and *PIF4*^S344D^ seedlings at 22℃ and 28℃ under SD conditions. Each bar represents mean ± s.d. of three independent experiments. *ARP6* transcript level at 28℃ was compared to its transcript level at 22℃ for each genotype. Student’s t test. ***; *p* value < 0.0005. **(I)** ChIP-qPCR showing the binding of PIF4, PIF4^S344A^ and PIF4^S344D^ proteins to the p*ARP6*-2 fragment in *pif4*, *PIF4-*3X*HA*, *PIF4*^S344A^ and *PIF4*^S344D^ seedlings grown at 22℃ and at 28℃ using anti-HA antibody. Each bar represents mean ± s.d. of three technical replicates. Significant enrichment of p*ARP6-2* fragment in *PIF4-3XHA*, *PIF4*^S344A^ and *PIF4*^S344D^ seedlings at 28℃ when compared to 22℃. Student’s t test. (*; *P* value <0.05, **; *P* value <0.005, ***; *p* value < 0.0005). Two independent experiments were performed with similar results.

Next, we studied the effect of MPK4 mediated PIF4 phosphorylation and *ARP6* repression on plant traits influenced by high temperature such as hypocotyl elongation and early flowering. Firstly, we examined the hypocotyl length phenotype of phosphor-null *PIF4*^S344A^-3X*HA* seedlings at 22 and 28℃ (Fig. 6, A D). We observed that *PIF4*^S344D^-3X*HA* seedlings showed longer hypocotyl length in comparison to *PIF4*-3X*HA* seedlings at 22℃ (Fig. 6, A and B), while the *PIF4*^S344A^-3X*HA* seedlings showed shorter hypocotyl length in comparison to *PIF4*-3X*HA* seedlings at 28℃ (Fig. 6, C and D). Then, we studied the flowering phenotype of phosphor-null *PIF4*^S344A^-3X*HA* and phosphor-mimetic *PIF4*^S344D^-3X*HA* seedlings at 22 and 28℃ (Fig. 6, E ^_^ H). *PIF4*^S314D^ plants showed early flowering in comparison to *PIF4*-3X*HA* plants at 22℃ (Fig. 6, E and F), while *PIF4*^S314A^ plants showed delayed flowering in comparison to *PIF4*-3X*HA* plants at 28℃ (Fig. 6, F and G). Taken together these findings strengthen the link between PIF4 phosphorylation, its regulatory role in *ARP6* expression, and the temperature-dependent modulation of plant traits like hypocotyl growth and flowering.

**Fig. 6.**
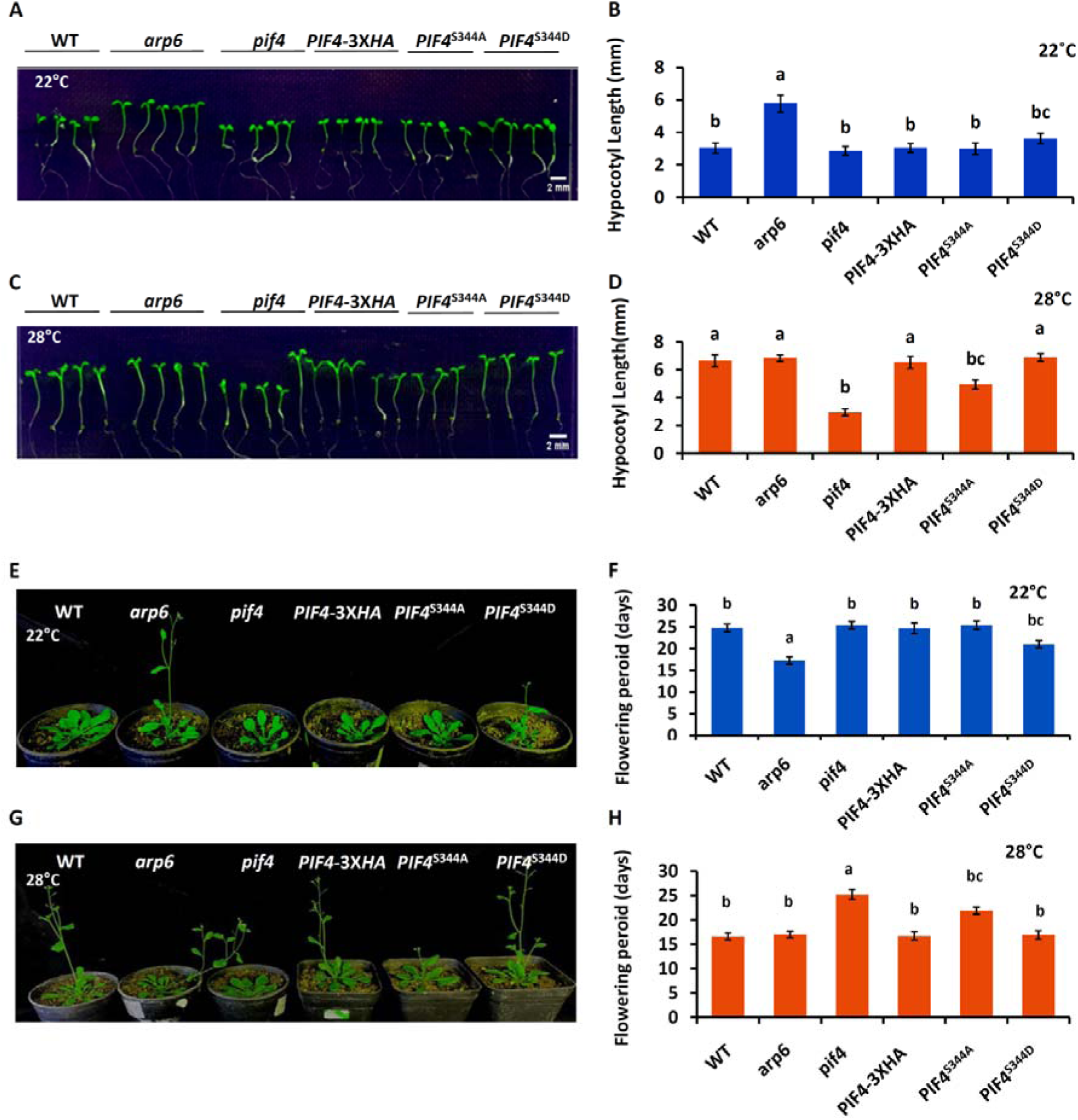
Effect of MPK4 mediated PIF4 phosphorylation on hypocotyl length and flowering S. **(A** and **B)** Hypocotyl phenotype (A) and length (B) comparison of WT, *arp6*, *pif4*, *PIF4-* 3X*HA*, *PIF4*^S344A^ and *PIF4*^S344D^ seedlings at 22 ℃. (B) Each bar represents mean ± s.d. (n = 20). Letters above the bars indicate significant differences (P < 0.05), as calculated by One-way ANOVA with Tukey’s post hoc analysis. **(C** and **D)** Hypocotyl phenotype (C) and length (D) comparison of WT, *arp6*, *pif4*, *PIF4-*3X*HA*, *PIF4*^S344A^ and *PIF4*^S344D^ seedlings at 28℃. (D) Each bar represents mean ± s.d. (n = 20). Letters above the bars indicate significant differences (P < 0.05), as calculated by One-way ANOVA with Tukey’s post hoc analysis. **(E)** Flowering phenotypes of WT, *arp6*, *pif4, PIF4-*3X*HA, PIF4^S344-A^* and *PIF4^S344-D^* plants at 22℃. Representative 22 days old *arp6* and *PIF4^S344-D^*plants showing early flowering in comparison to WT, *pif4*, *PIF4-*3X*HA* and *PIF4^S344-A^* plants at 22℃ under SD conditions. **(F)** Comparison of flowering time among WT, *arp6*, *pif4, PIF4-*3X*HA, PIF4^S344-A^* and *PIF4^S344-D^* plants. **(G)** Flowering phenotypes of WT, *arp6*, *pif4, PIF4-*3X*HA, PIF4^S344-A^* and *PIF4^S344-D^*plants 28℃. Representative 22 days old *pif4* and *PIF4^S344-A^* plants showing delayed flowering in comparison to WT, *arp6*, *PIF4-*3X*HA,* and *PIF4^S344-D^* plants at 28℃ under SD conditions. **(H)** Comparison of flowering time among WT, *arp6*, *pif4, PIF4-*3X*HA, PIF4^S344-A^* and *PIF4^S344-D^* plants. (F and H) The days of flowering was calculated as the number of days from germination to bolting at 22℃ and 28℃ under SD conditions. Letters above the bars indicate significant differences (P < 0.05), as calculated by One-way ANOVA with Tukey’s post hoc analysis.

### *ARP6* repression by phosphorylated PIF4 at elevated temperature increases hypocotyl cell length by eviction of H2A.Z from genes involved in cell elongation

To explore how *ARP6* repression byMPK4 mediated PIF4 phosphorylation at elevated temperature increases hypocotyl length, we first measured the length of epidermal cells in the middle portion of hypocotyl between shoot apical meristem and collet in WT, *arp6*, *pif4*, *PIF4*-3X*HA*, phosphor-null *PIF4*^S344A^-3X*HA* and phosphor-mimetic *PIF4*^S344D^-3X*HA* seedlings at 22 and 28℃. A detailed analysis of confocal microscopy images revealed that hypocotyl cell length in WT, *pif4*, *PIF4*-3X*HA* and phosphor-null *PIF4*^S344A^-3X*HA* seedlings was shorter than in *arp6* and phosphor-mimetic *PIF4*^S344D^-3X*HA* seedlings at 22℃ (Fig. 7, A and B). In the case of an elevated temperature of 28℃, *pif4* and phosphor-null *PIF4*^S344A^-3X*HA* seedlings showed shorter hypocotyl cells length in comparison to WT, *arp6*, *PIF4*-3X*HA* and phosphor-mimetic *PIF4*^S344D^-3X*HA* seedlings (Fig. 7, C and D). A deeper analysis of the confocal microscopy images revealed that the numbers of cells in the hypocotyls were highly similar among all genotypes at 22 and 28℃ (Supplemental Fig. S7, A and B). This analysis showed that PIF4 phosphorylation is responsible to increase the hypocotyl cell length as in the case of phosphor-mimetic *PIF4*^S344D^-3X*HA* seedlings, hypocotyl cells length is similar to that of *arp6* seedlings even at a low ambient temperature of 22℃. While no phosphorylation of PIF4 at an elevated temperature of 28℃ in the case of phosphor-null *PIF4*^S344D^-3X*HA* seedlings results in a shorter hypocotyl length similar to that of *pif4.* To further elaborate our understanding of the increase of hypocotyl length by PIF4 phosphorylation-mediated *ARP6* repression at 28℃, we performed ChIP experiment to examine the H2A.Z occupancy at loci of genes involved in cell elongation at 22 and 28℃ in WT, *arp6*, *pif4*, *PIF4*-3X*HA*, phosphor-null *PIF4*^S344A^-3X*HA*and phosphor-mimetic *PIF4*^S344D^-3X*HA* seedlings. For this experiment, we selected *EXP2*, *IAA19*, and *XTH33* genes that are reported to be known targets of H2A.Z occupancy and promote cell elongation in *arp6* mutant (Mao et al., 2021). As demonstrated earlier that the phosphor-null PIF4^S344A^ protein is not phosphorylated by MPK4 (Fig. 4, C and E) and *ARP6* expression increases in *PIF4*^S344A^-3X*HA* seedlings at 28℃ (Fig. 5H), an increased *ARP6* expression might promote H2A.Z deposition at the loci of *EXP2*, *IAA19*, and *XTH33* genes and inhibit hypocotyl elongation in *PIF4*^S344A^-3X*HA* seedlings at 28℃. H2A.Z containing chromatin was immunoprecipitated using anti-H2A.Z antibodies. Successful immunoprecipitation was examined by immunoblotting of H2A.Z from immunoprecipitated chromatin using anti-H2A.Z antibodies (Fig. 7E). Comparing WT *PIF4*-3XHA complemented seedlings at 22 and 28℃, we observed more H2A.Z deposition at the *EXP2*, *IAA19*, and *XTH33* genes at 22℃, whereas at the elevated temperature of 28℃, H2A.Z depositions were significantly reduced (Fig. 7F). ChIP experiment showed an increase in H2A.Z occupancies at *EXP2*, *IAA19*, and *XTH33* genes at 28℃ in *PIF4*^S344A^-3X*HA* seedlings in comparison to *PIF4*-3X*HA* seedlings and similar to that of *pif4* mutant (Fig. 7F). In addition to that, *PIF4*^S344D^-3X*HA* seedlings showed a decrease in H2A.Z insertion at *EXP2*, *IAA19*, and *XTH33* genes at 22℃ in comparison to *PIF4*-3X*HA* seedlings similar to that of *ARP6* mutant (Fig. 7F). To further validate these observations, we examined the expression of *EXP2*, *IAA19*, and *XTH33* genes in *PIF4*-3X*HA*, *PIF4*^S344A^-3X*HA* and *PIF4*^S344D^-3X*HA* seedlings at 22 and 28℃. Transcripts level of these genes were found to be reduced in *PIF4*^S344A^-3X*HA* seedlings in comparison to *PIF4*-3X*HA* seedlings at 28℃ (Supplemental Fig. S7, C ^_^ E). In case of *PIF4*^S344D^-3X*HA* seedlings, *EXP2*, *IAA19*, and *XTH33* genes showed an increase in expression in comparison to *PIF4*-3X*HA* seedlings at 22℃ (Supplemental Fig. S7, C ^_^ E). Collectively, these data confirm that *ARP6* repression by MPK4-mediated phosphorylation of PIF4 at elevated temperature increases hypocotyl length by the eviction of H2A.Z from the loci of genes involved in cell elongation.

**Fig. 7.**
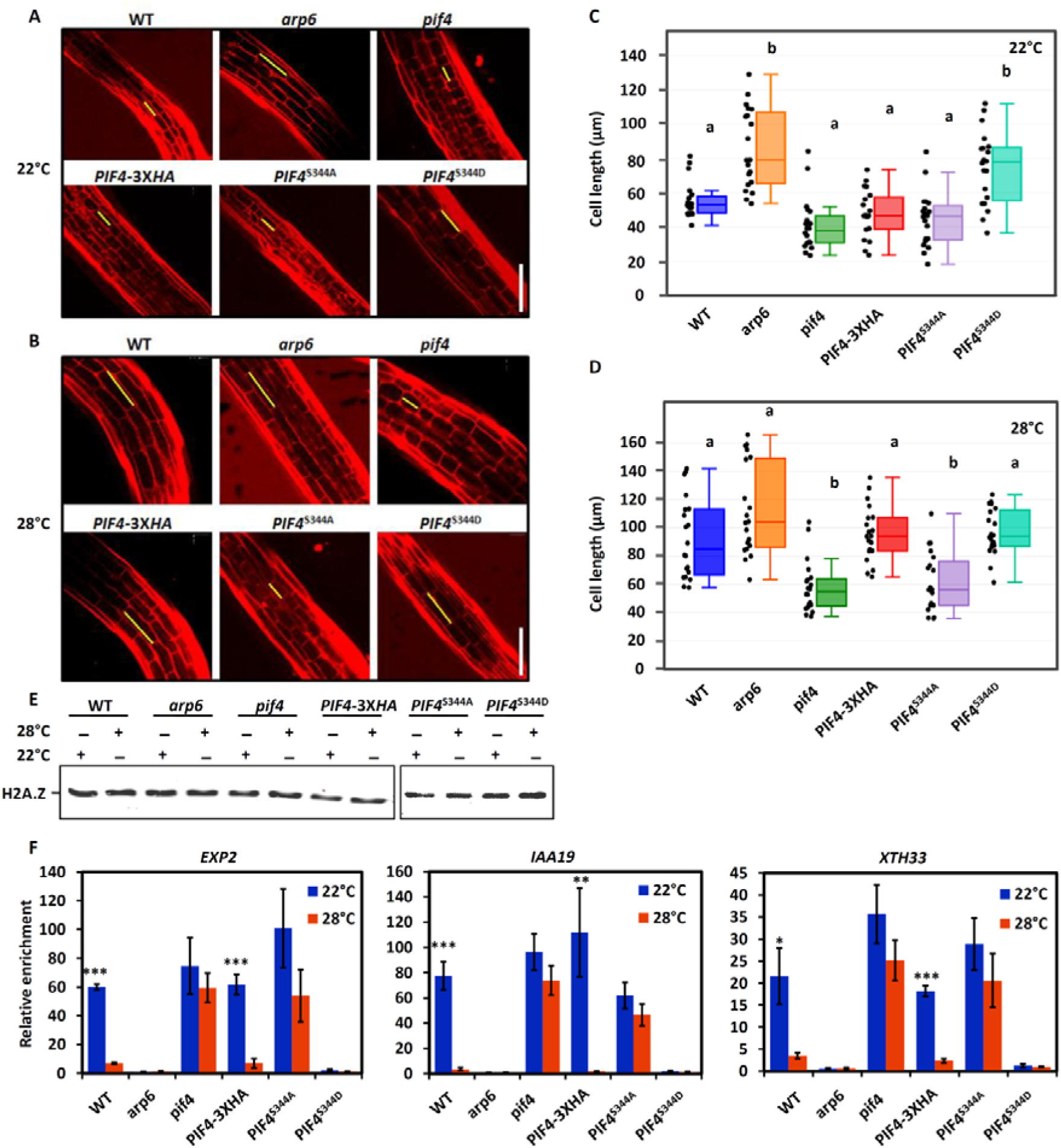
Role of PIF4 phosphorylation in hypocotyl elongation by the eviction of H2A.Z from genes involved in cell elongation. **(A and B)** PI staining of hypocotyls in the 10 days old WT, *arp6*, *pif4*, *PIF4-*3X*HA*, phosphor-null *PIF4*^S344A^ and phosphor-mimic *PIF4*^S344D^ seedlings at 22℃ (A) and at 28℃ (B) The yellow line represents the length of a single cell. Scale bar, 150 μm. **(C and D)** Whisker and box plots show the mean cell length of the non-dividing cells of hypocotyls in 10 days old WT, *arp6*, *pif4*, *PIF4-*3X*HA*, *PIF4*^S344A^ and *PIF4*^S344D^ seedlings at 22℃ (C) and at 28℃ (D). Significant differences (*P* < 0.05) among genotypes were calculated using ANOVA followed by Tukey’s posthoc analysis. (n=20 cells length were measured for two hypocotyls of each genotype). **(E and F)** ChIP-qPCR showing the role of PIF4 phosphorylation mediated *ARP6* repression in H2A.Z eviction at the loci of *EXP2*, *IAA19*, and *XTH33* genes involved in hypocotyl cell elongation at 28℃. Chromatin of 10 days old WT, *arp6*, *pif4*, *PIF4-*3X*HA*, *PIF4*^S344A^ and *PIF4*^S344D^ seedlings grown at 22℃ and 28℃, were immunoprecipitated using anti H2A.Z antibodies. (E) Immunoprecipitation of H2A.Z containing chromatin was tested by immunoblotting using anti-H2A.Z antibodies. (F) Insertion of H2A.Z at *EXP2*, *IAA19*, and *XTH33* gene loci was analyzed by qPCR using the immunoprecipitated chromatin with the primer pairs designed around the +1 nucleosome of these genes. Bar graphs showing the enrichment of *EXP2*, *IAA19* and *XTH33* relative to that of *Actin.* Each bar represents the mean ± s.d. of three technical replicates. H2A.Z occupancy at 22℃ was compared to 28℃. Student’s t test (*; *P* value <0.05, **; *P* value <0.005, ***; *P* value <0.0005). Two independent experiments were performed with similar results.

### Temperature dependent H2A.Z deposition at *MPK4* locus regulates its expression

Next, we show that the expression of *MPK4* is repressed by H2A.Z deposition at an ambient temperature of 22℃. The expression pattern of *MPK4* and its protein level in WT at ambient and elevated temperatures as shown earlier, are similar to that of thermoresponsive genes such as *HSP70* which is reported to be induced at high temperature and repressed at ambient temperature by H2A.Z deposition (Petesch & Lis, 2008; Kumar & Wigge, 2010; Cortijo et al., 2017). Therefore we established a notion of regulation of *MPK4* expression by H2A.Z deposition. To elucidate this, we analyzed the expression of *MPK4* in the *arp6* mutant. Interestingly in comparison to the WT, *arp6* mutant showed elevated expression of *MPK4* at ambient temperature of 22℃ (Fig. 8A). At the protein level also, *arp6* seedlings showed increased level of MPK4 protein at 22℃ in comparison to WT seedlings (Fig. 8B). These observations further supported our notion of regulation of *MPK4* expression by H2A.Z deposition. Therefore, ChIP experiment was performed to examine the H2A.Z occupancy at *MPK4* gene locus using 10 days old seedlings of WT & *arp6* grown at 22℃ and 28℃. The histone variant H2A.Z containing chromatin was immunoprecipitated from these seedlings using anti-H2A.Z antibody. The presence of *MPK4* in the immunoprecipitated H2A.Z containing chromatin was analyzed by qPCR using seven combinations of MPK4 oligos (*MPK4*-1-*MPK4*-7) which were designed from -184 to +330 bp with respect to transcription start site (TSS) (Fig. 8C). The results of ChIP-qPCR analysis are shown in Fig. 8D. *MPK4*-4 fragment from +48 bp to +144 bp including TSS, showed highest enrichment of H2A.Z in WT seedlings at 22℃. *MPK4*-3 & *MPK4*-5 fragments also showed H2A.Z enrichment but lesser than that of *MPK4*-4 in WT seedlings at 22℃ while no H2A.Z enrichment was observed in case of *MPK4*-1, *MPK4*-2, *MPK4*-6 & *MPK4*-7 fragments. H2A.Z occupancy at *MPK4*-3, *MPK4*-4 & *MPK4*-5 fragments reduced drastically in WT seedlings at 28℃. No enrichment of H2A.Z was observed in case of *arp6* mutant seedlings. These results implicate H2A.Z deposition at the *MPK4* locus to the repression of *MPK4* gene expression at 22℃.

**Fig. 8.**
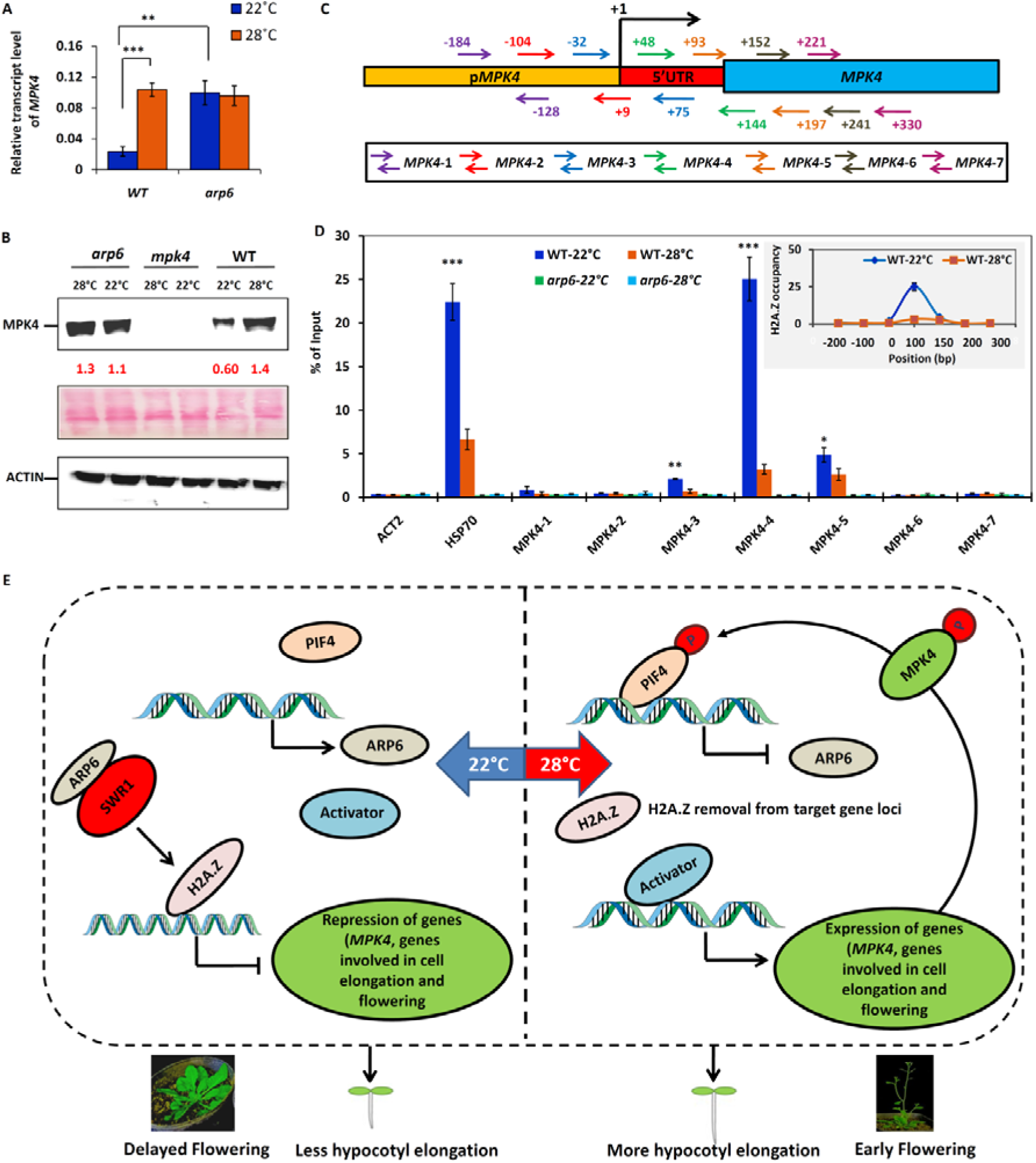
*MPK4* expression is regulated by ARP6 mediated deposition of H2A.Z in temperature dependent manner. **(A)** *MPK4* transcript in 10 days old WT and *arp6* seedlings grown at 22℃ and 28℃ under SD. Each bar represents mean ± s.d. of three independent experiments. *MPK4* Expression in WT seedlings at 22℃ was compared to its expression in WT at 28℃ and *arp6* at 22℃. (n= 3) Student’s t test: **; *p* value < 0.005; ***; *p* value < 0.0005. **(B)** Immunoblotting of MPK4 in 10 days old WT, *arp6* & *mpk4* seedlings grown at 22 and 28℃. Ponceau staining of rubisco was used as loading control. Numbers in red indicate the intensity of MPK4 protein bands relative to Actin bands. Three independent experiments were performed with similar results. **(C)** Schematic representation of positions (with respect to TSS) of 7 pairs of *MPK4* oligos (colour coded) used in ChIP-qPCR analysis of H2A.Z deposition at *MPK4* locus. **(D)** ChIP-qPCR assays performed in 10 days old WT and *arp6* seedlings grown at 22 and 28℃ using anti-H2A.Z antibody to detect H2A.Z deposition at the *MPK4* locus. DNA fragments eluted from immunoprecipitated complex, were analyzed quantitatively using *MPK4* oligos. ChIP-qPCR of *HSP70* was used as a positive control. Each bar represents mean ± s.d. of three technical replicates. H2A.Z occupancy at 22℃ was compared to 28℃. Student’s t test (*; *P* value <0.05, **; *P* value <0.005, ***; *P* value <0.0005). Inset line graph shows H2A.Z occupancy at MPK4 locus with respect to TSS in WT at 22 and 28℃. Two independent experiments were performed with similar results. **(E)** Model showing the interplay of MPK4, PIF4 and ARP6/H2A.Z in the regulation of thermosensing. Elevated temperature induces *MPK4* expression and MPK4 kinase activity. Activated MPK4 phosphorylates PIF4, which represses *ARP6* expression resulting in H2A.Z depletion at its target gene loci. H2A.Z depletion increases the expression of *MPK4* and the genes involved in hypocotyl cell elongation and flowering.

## Discussion

PIF4 TF plays a crucial role in plant’s response to elevated temperature and is necessary for the induction of the thermosensing pathway that initiates morphological changes to instigate plant survival. PIF4 is known to interact with multiple proteins in response to various environmental stimuli modulating its activity and influencing the expression of its target genes to facilitate the plant’s adaptation to a specific condition. In this study, we demonstrate the significance of *MPK4*-mediated phosphorylation of *PIF4* at elevated temperature in repressing the expression of the *ARP6* gene, ultimately leading to the depletion of H2A.Z from the genes involved in hypocotyl elongation. Furthermore, we uncover a reciprocal relationship, where the expression of *MPK4* is influenced by the ARP6/SWR1-mediated insertion of H2A.Z in a temperature-dependent manner. These findings provide insights into the intricate thermosensory regulatory network involving PIF4, MPK4, ARP6, and H2A.Z, highlighting their interconnected roles in temperature dependent plant adaptation.

PIF4 has been extensively studied for its role in promoting hypocotyl elongation in Arabidopsis seedlings in response to high temperatures (Koini et al., 2009; Kumar et al., 2012; Gangappa & Kumar, 2017; Oh et al., 2012; Proveniers et al., 2013; Gil and Park, 2019; Zhou et al., 2023). Consistent with these findings, our study also revealed that *pif4* mutant seedlings exhibited reduced hypocotyl length specifically at the elevated temperature of 28℃ compared to wild-type (WT) seedlings. Interestingly, we observed an opposite effect in *arp6* mutant seedlings which showed increased hypocotyl length compared to WT at an ambient temperature of 22℃. Notably, the hypocotyl length of *arp6* seedlings at 22℃ closely resembled that of WT seedlings at 28℃, highlighting the contrasting patterns of hypocotyl elongation between *pif4* and *arp6* mutants at different temperatures. These observations prompted us to investigate a potential relationship between PIF4 and ARP6 in the thermosensing mechanism. Through gene expression analysis and promoter-reporter gene (p*ARP6*-*GUS*) assays, we found compelling evidence that PIF4 TF actively represses the expression of *ARP6* specifically in seedlings grown at the elevated temperature of 28℃ (Fig. 1). The results from expression analysis of *ARP6* in the *pif4* mutant plants and GUS assay led us to hypothesize that the PIF4 transcription factor potentially interacts with the *ARP6* promoter to function as a transcriptional repressor. In order to test and validate the hypothesis, our initial approach was to conduct the *in silico* docking of PIF4 protein at *ARP6* promoter. Like many other bHLH transcription factors, PIF4 functions by binding to DNA as both a homodimer and a heterodimer (Ezer et al., 2014; Shively et al., 2019). Therefore, we conducted the docking experiment using a homodimer of PIF4 which showed binding to both the G-boxes present in the promoter of *ARP6*. Further, EMSA and ChIP-qPCR assays showed the direct binding of PIF4 at *ARP6* promoter (Fig. 2). These results suggest that PIF4 and ARP6 play opposite roles in thermosensing mechanism, working together to integrate the responses of temperature transcriptome to regulate the thermoresponsive growth of plants.

Given that PIF4 acts as a repressor of *ARP6* expression specifically at elevated temperature as observed in case of *PIF4*-OE seedlings (Fig. 1F), we explored the potential involvement of PIF4 interacting proteins in modulating its activity in this context. Previous studies have highlighted phosphorylation as a prevalent post-translational modification of PIF4 protein (Lorrain et al., 2008; Christians et al., 2012; Oh et al., 2012; Ni et al., 2014; Xu et al., 2015; Bernardo-Garcia et al., 2014; Lee et al., 2020). In light signaling, the active form of phyB, Pfr, phosphorylates PIF4, leading to its degradation through the 26S proteasome pathway (Lorrain et al., 2008; Christians et al., 2012). Similarly, in the brassinosteroid hormone signaling pathway, the BR-regulated kinase BIN2 interacts with and phosphorylates PIF4, triggering its degradation via the 26S proteasome (Bernardo-Garcia et al., 2014). These phosphorylation events induced by light and BR signaling reduce PIF4 stability. However, it has been shown that temperature induced phosphorylation of PIF4 promotes its stability (Foreman et al., 2011; Lee et al., 2020; Han et al., 2022). A recent report has shown that the SUPPRESSOR OF PHYA-105 (SPA) protein induces phosphorylation of PIF4 at high temperatures, contributing to its increased stability (Lee et al., 2020). In our study, potential phosphorylating partner of PIF4 was identified by monitoring the expression of MAP kinase genes in the seedlings grown at elevated temperature. Upregulated *MPK4* also showed kinase activity at elevated temperature and was found to interact with PIF4 (Fig. 3). Andrasi et al. (2019) implicated the role of MPK4 in phosphorylation of HSFA4A in Arabidopsis seedlings under heat stress. In our study, MPK4 directly phosphorylates PIF4 *in vitro* and *in vivo* at elevated temperature of 28℃ (Fig. 4), and phosphorylation of PIF4 is necessary for the repression of *ARP6* expression (Fig. 5). *ARP6* repression is observed only when *MPK4* and *PIF4* both are transiently expressed in *pif4/mpk4* double mutant seedlings at 28℃. In addition to that, *ARP6* expression was found to be increased at 28℃ in phosphor-null *PIF4*^S344A^ plants when compared to *PIF4*-3X*HA* plants. Moreover, MPK4-driven phosphorylation of PIF4 does not affect the stability of PIF4 protein as *PIF4*-3X*HA* and *PIF4*^S344A^ plants showed similar levels of PIF4 and PIF4^S344A^ proteins, respectively at 28℃ (Fig. 4D). Here we show that MPK4-mediated phosphorylation of PIF4 is essential for thermosensing through *ARP6* repression at elevated temperatures. Thus, different kinases function differently to fine tune the PIF4 activity ensuring precise regulation of plant responses to varying environmental conditions.

Being an important subunit of SWR1 complex, ARP6 plays an important role in H2A.Z insertion at its target gene loci (Mizuguchi et al., 2004; Cortijo et al., 2017; Dai et al., 2017; Mao et al., 2021). Previous studies reported a dynamic increase in H2A.Z eviction at +1 nucleosomes of high temperature-induced genes in WT with increasing temperature and *arp6* mutant displays a constitutive warm temperature response by the derepression of thermoresponsive genes such as *HSP70*, *DREB2A* and *YUC8* even at ambient temperature (Kumar & Wigge, 2010; Cortijo et al., 2017; van der Woude et al., 2019). The expression of *FT* gene is also controlled by H2A.Z insertion in temperature dependent manner (Zilberman et al., 2008; Kumar & Wigge, 2010). *FT* gene plays an important role in thermosensory flowering pathway and induces early flowering at elevated temperature (Balasubramanian et al., 2006; Kumar et al., 2012). Interestingly, H2A.Z is released from *FT* gene locus at high temperature allowing PIF4 TF to bind to the *FT* gene promoter, which accelerates the expression of *FT* gene leading to early flowering transition (Kumar & Wigge. 2010; Kumar et al., 2012). In our study, *PIF4*^S314D^ plants showed early flowering at 22℃ similar to that of *arp6* plants, while *PIF4*^S314A^ plants showed delayed flowering at 28℃ similar to *pif4* plants (Fig. 6). Thus, we suggest that *ARP6* repression by PIF4 phosphorylation at 28℃ may result into H2A.Z depletion at *FT* gene locus leading to early flowering. Our study also explored the molecular mechanism involved in hypocotyl cell elongation at elevated temperature of 28℃. Analysis of hypocotyl cell length measurement in phosphor-null *PIF4*^S344A^ and phosphor-mimic *PIF4*^S344D^ seedlings revealed that phosphorylation of PIF4 is necessary for the expansion of hypocotyl cell length at elevated temperature. H2A.Z depletion due to *ARP6* repression by phosphorylated PIF4 leads to derepression of genes involved in the expansion of hypocotyl cell length at elevated temperature (Fig. 7). Thus, our study adds an interesting information to this PIF4 and H2A.Z mediated thermosensory regulatory network that MPK4 mediated phosphorylation of PIF4 represses the expression of *ARP6* at elevated temperature, which is the reason behind the depletion of H2A.Z from the target gene loci involved in hypocotyl elongation and early flowering phenotypes.

Because MPK4 was found to play an essential role in PIF4 mediated thermosensory pathway and its expression is influenced by temperature, we were keen to analyze if *MPK4* expression is controlled by H2A.Z deposition at its gene locus. Investigation of *MPK4* gene locus for H2A.Z occupancy by ChIP revealed that H2A.Z is enriched at *MPK4* locus at an ambient temperature of 22℃ and depleted at elevated temperature of 28℃ (Fig. 8). This finding leads us to conclude that *MPK4* expression is controlled by the transcriptional repressive effect of H2A.Z deposition. Strikingly, the deposition of H2A.Z at *MPK4* locus was observed around TSS, which is in agreement with the previous reports affirming that the H2A.Z occupancy around TSS at +1 nucleosome represses the gene expression (Kumar & Wigge, 2010; Dai et al., 2017). H2A.Z depletion at *MPK4* locus at 28℃ might be a consequence of PIF4 mediated repression of *ARP6* at elevated temperature. Together, these findings suggest a feedback regulation of *MPK4* by ARP6 mediated deposition of H2A.Z at ambient temperature of 22℃ and release of H2A.Z due to *ARP6* repression at elevated temperature of 28℃.

In summary, based on our findings, we propose a model which shows an interplay among PIF4, ARP6 and MPK4 to regulate thermosensing mechanism (Fig. 8E). When Arabidopsis plants sense elevated temperature of 28℃, MPK4 mediated phosphorylation of PIF4 causes the repression of *ARP6* leading to the removal of H2A.Z from its target gene loci and more hypocotyl elongation with early floering. While at an ambient temperature, PIF4 is not phosphorylated due to less expression and no activation of kinase activity of MPK4 protein resulting in the *ARP6* expression which causes deposition of H2A.Z at its target gene loci and transcriptional repression of these genes leading to less hypocotyl elongation and delayed flowering.

## Materials & Methods

### Plant material, screening of mutants and phenotypic analysis

All the seeds used in this study were in Columbia-0 (Col-0) ecotype background. *pif4* (SAIL_1288_E07)*, arp6* (SALK_037471C) & *mpk4* (SALK_056245) mutant seeds were procured from the Arabidopsis Biological Resource Centre (ABRC) of Ohio State University (Columbus, USA). The positions of TDNA integration in the gene bodies of these mutants have been depicted in Supplemental Fig. S8A. *pif4* and *arp6* seeds were found to be homozygous mutants as confirmed by the genotyping PCR (Supplemental Fig. S8B). *mpk4* seeds procured from ABRC were heterozygous (Supplemental Fig. S8C) and hence heterozygous *mpk4* plants were self-pollinated to produce homozygous *mpk4* mutant which were screened for the homozygosity of *mpk4* mutation by PCR using allele specific primers (Supplemental Fig. S9D). The primer sequences for the screening of *pif4*, *arp6* and *mpk4* mutants are shown in Supplemental Table S1. Homozygous *mpk4* mutant plants produced very few seeds. Seeds of *PIF4*-OE line (CS69173) were also procured from ABRC. Seeds were surface sterilized and germinated on 1/2 MS media. Seeds stratification was carried out by putting them at 4 ℃ for 3 days. Then the seeds were placed at 22 ℃ for 2 days and then transferred to the required temperature of 18℃, 22℃, 25℃ and 28℃ for 8 more days under SD conditions for expression analysis. For hypocotyl length measurement, the seeds were transferred to 22℃ or 28℃ for more 8 days under SD conditions. Hypocotyl length was measured using 20 seedlings of each genotype. The flowering phenotype was analysed according to the previous study^40^. Seeds were directly sown in the potted soil and plants were grown under SD condition and flowering time was monitored as the number of days from germination till bolting. 10 plants of each genotype were examined.

### Generation of transgenic plants

p*ARP6-GUS* transgenic plants were generated in Arabidopsis WT and *pif4* mutant background by infiltration of *Agrobacterium* strain GV3101 harboring p*ARP6-GUS* construct into buds by applying vacuum. Homozygous p*ARP6-GUS* F2 transgenic lines in WT and *pif4* background were screened by putting ∼1000 F3 seeds of 10 F2 plants on kanamycin selection. The F2 lines with all 1000 F3 seeds showing kanamycin resistance were used for studying *ARP6* promoter activity by GUS histochemical staining and quantitative measurement. Similarly, p*MPK4-GUS* transgenic lines were developed in WT plants. *PIF4*-3X*HA* complementation lines were generated by the genetic transformation of *pif4* mutant plants with *Agrobacterium* strain GV3101 harboring p*PIF4*: *PIF4*-3X*HA* construct. Schematic representation of this construct is shown in Supplemental Fig. S9A. Phosphor-null *PIF4*^S344A^-3X*HA* transgenic plants were generated by the genetic transformation of *pif4* mutant plants with *Agrobacterium* strain GV3101 harboring p*PIF4*: *PIF4*^S344A^-3X*HA* construct (Supplemental Fig. S9A). Phosphor-null version (*PIF4*^S344A^) of *PIF4* was generated by replacing serine-344 with alanine. Phosphor-mimic *PIF4*^S344D^-3X*HA* transgenic plants were generated by the genetic transformation of *pif4* mutant plants with *Agrobacterium* strain GV3101 harboring p*PIF4*: *PIF4* ^S344D^-3X*HA* construct (Supplemental Fig. S9A). Phosphor-mimic version (*PIF4* ^S344D^) of *PIF4* was generated by replacing serine-344 with aspartic acid. Transgenic lines were screened by selection of F1 seeds on kanamycin containing medium. Presenece of *nptII* gene was confirmed in the kanamycin resistant F1 lines of these transgenics (Supplemental Fig. S9, B and C). Homozygous *PIF4*-3X*HA*, *PIF4*^S344A^-3X*HA* and *PIF4*^S344D^-3X*HA* complemented F2 lines were screened by kanamycin resistance as described earlier. Transcript and protein levels of PIF4, PIF4^S344A^ and PIF4^S344D^ in these transgenic lines were confirmed by qPCR and immunoblotting using anti-HA antibody, respectively (Supplemental Fig. S10, A ^_^ D).

### Gene expression analysis

Total RNA was isolated from 100 mg Arabidopsis seedlings using RNeasy Plant Mini Kit (Qiagen). DNase treatment and cDNA synthesis was carried out using QuantiTect Reverse Transcription Kit (Qiagen). Expression of *PIF4* and *ARP6* was analyzed by semiqunatitative PCR and qPCR, and the expression of MAP kinase genes was analyzed by qPCR. The conditions for semi quantitative PCR were as follows 95℃ for 2 min; 25 cycles of 95℃ for 30 s; primer annealing for 30 s; 72℃ for 30 s; one cycle at 72℃ for 7 min for *PIF4* and 95℃ for 2 min; 30 cycles of 95℃ for 30 s; primer annealing for 30 s 72℃ for 30 s; one cycle at 72℃ for 7 min for *ARP6*. Quantitative PCR was performed using a *Power* SYBR green PCR master mix by Applied Biosystem with the following protocol: 40 cycles at 95 ℃ for 15 s, 60℃ for 60 s. *Actin* was used as internal reference gene for the normalization of relative transcript level (Czechowski et al., 2005). The primers for quantitative RT-PCR are listed in Supplemental Table S2.

### Histochemical GUS staining

Histochemical GUS staining was performed as described previously previously (Jefferson et al., 1987; De Block and De Brouwer, 1992). 10 days old transgenic seedlings grown at 22℃ & 28℃, were harvested and vacuum infiltrated in the reaction solution containing 1mM 5-bromo-4-chloro-3-indolyl-b-D-glucuronic acid, 50mM sodium phosphate, 1mM ferricyanide, 1mM ferrocyanide and 0.1% Triton X-100, pH 7.0. The seedlings were then incubated overnight at 37℃ in the dark. Chlorophyll of stained seedlings was removed by a series of ethanol washes with decreasing concentration (from 90% to 50%) for 30 min each and pictures of the stained seedlings were taken.

### Quantification of GUS activity

Total protein from the seedlings was extracted in GUS extraction buffer and quantified using Bradford method (Bradford, 1976). GUS fluorescence was measured using 4-methylumberrifyl-β-glucuronide (MUG) substrate as described previously (Jefferson et al., 1987). The product 4-methylumbelliferone (MU) of reaction was quantitated at excitation wavelength of 360 nm and emission wavelength of 460 nm. GUS activity was measured at two time points (after 10 and 20 minutes) for each sample. GUS activity was calculated as pmolMU/min/ mg protein.

### *In silico* docking of PIF4 TF to *ARP6* promoter

The 3D structure of PIF4 and its bHLH domain were predicted using AlphaFold. The PIF4 bHLH domain dimer was obtained by *in-silico* protein-protein docking using the HDock server. The docking of the bHLH dimer to *ARP6* promoter DNA was performed using the HDock server. The docked structures were analysed using the PyMoL standalone software.

### Yeast one-hybrid (Y1H) assay

ARP6 promoter fragments were cloned upstream to Aureobasidin resistance reporter gene *AbA^r^*into bait vector pAbAi. pAbAi-*pARP6-1*, pAbAi-*pARP6-2 &*pAbAi-*pARP6-3* plasmids were linearized with BstBI, and transformed into Y1H Gold yeast strain, respectively. Transformants were grown on SD/-Ura medium and integration of *ARP6* promoter fragments into the yeast genome was confirmed by PCR. PIF4 TF was cloned into prey vector pGADT7. pGADT7-*PIF4* plasmid was transformed into the *pARP6*-specific reporter strain, respectively. Transformants were grown on SD/-Leu medium. Interaction of PIF4 TF with the *ARP6* promoter was confirmed by growing the transformants on SD/-Leu/-AbA medium.

### Electrophoretic mobility sift assay (EMSA)

The DNA-protein interaction between *ARP6* promoter and PIF4 transcription factor was performed by EMSA. Two DNA probes (-462 to -416 bp and -735 to -686 bp) containing the G box elements were designed radiolabelled and used for the binding experiment with bacterially purified WT PIF4 and phosphor-null PIF4^S344A^ proteins as described previously (Singh and Sinha, 2016). Briefly, radiolabelled probes were either incubated alone or with protein in the reaction buffer (20mM HEPES, pH 7.4; 10mM KCl; 0.5mM EDTA; 0.5mM DTT; 1mM MgCl_2_; 3% Glycerol and 1 μg of poly(dI-dC) at room temperature for 30 minutes. The DNA-protein complex was separated on 5% PAGE using 0.5x TBE running buffer and auto-radiographed using typhoon.

### Yeast two-hybrid (Y2H) assay

The interaction between PIF4 and MPK4 was examined using the Matchmaker gold yeast two-hybrid system (Takara Bio). *PIF4* and *MPK4* cDNA were cloned in the pGADT7 and pGBKT7 vectors. The yeast transformation was performed according to the Fast yeast transformation kit (Gbioscience: GZ-1). After mating, the transformants were grown on SD/-Ade-His-Leu-Trp dropout plates.

### Bimolecular Fluorescence Complementation assay (BiFC)

For the BiFC assay, cDNA sequences of *MPK4* and *PIF4* were cloned into the pSITE-nEYFP-C1 (CD3-1648) and pSITE-cEYFP-N1(CD3-1651) vectors, respectively to generate YFP^N^-*MPK4* and *PIF4*-YFP^C^constructs. The cloned constructs were transformed into EHA105 *Agrobacterium* cells. *Agrobacterium* cells containing these constructs were then used to infiltrate the *Nicotiana benthamiana* leaves. Interaction was analyzed after 48-72 hours of infiltration using Leica TCS SP5 AOBS Laser Scanning Microscope (Leica Microsystem) using YFP and mCherry filter. The fluorescence excitation at 514nm and an emission at 527nm were used.

### Chromatin immunoprecipitation (ChIP) assays

Seedlings of WT, *pif4, arp6*, *PIF4*-3X*HA*, *PIF4* ^S344A^-3X*HA* and *PIF4* ^S344D^-3X*HA* were grown at 22℃ and 28℃ under SD conditions for 10 days and then harvested. Samples were crosslinked using1% formaldehyde by applying vacuum for 15 min followed by the quenching with 2M glycine for 5 min. Samples were rinsed with 10 ml cold 1x PBS solution for three times and dried by blotting on paper towel. Then, the dried samples were ground in liquid nitrogen. Ground samples were added into nuclei extraction buffer (100 mM MOPS pH 7.6, 10 mM MgCl2, 0.25 M Sucrose, 5 % Dextran T-40, 2.5 % Ficoll 400, 40 mM b-Mercaptoethanol& 1 × Protease Inhibitor) for 15 min and then centrifuged at 20,200 × g for 10 min at 4 ℃. Pellets were resuspended into nuclei lysis buffer (50 mM Tris-HCl pH 8.0, 10mM EDTA pH 8.0, 1 % SDS & 1 × Protease Inhibitor) and put on ice for 30 min followed by adding ChIP dilution buffer without Triton. Then in case of analyzing PIF4 TF binding to *ARP6* promoter, chromatin of *pif4*, *PIF4*-3X*HA*, *PIF4* ^S344A^-3X*HA*, and *PIF4* ^S344D^-3X*HA* was sonicated five times for 10 seconds each (∼ P3.2) followed by addition of ChIP dilution buffer with Triton X-100 and 22 % Triton X-100 to attain a final 1.1 % Triton X-100 concentration. ChIP grade anti-HA antibody (Abcam) was used for immunoprecipitation of PIF4 bound chromatin in the extract of *pif4* and *PIF4*-3XHA seedlings. To analyze H2A.Z insertion at its target gene loci, chromatin of respective genotype was fragmented using 0.2 units of MNase for 20 min. Digestion was terminated by adding 5mM EDTA. Anti-H2A.Z antibody (Abcam; ab4174;) was used for immunoprecipitation of chromatin having H2A.Z deposition. Immunoprecipitated DNA/ protein complex was captured using magnetic beads. The beads were washed with low salt buffer, high salt buffer, LiCL wash buffer and TE buffer. Elution was carried out with elution buffer and the eluted samples were mixed with NaCl with final 0.2M and incubated overnight at 65℃ for reverse crosslinking. Next day, samples were given proteinase K treatment for 1 hour and DNA was purified using Qiagen PCR purification kit. Samples with no immunoprecipitation were used as input control DNA. % of input was calculated using the following formula. Percent of Input =100×2ΔCt ΔCt= [Ct^Input^-log_2_(Input dilution factor)]-Ct^ChIP^ The sequences of primers used in ChIP qPCR is presented in Supplemental Table S3.

### Protein extraction and immunoblotting

Total protein was extracted from 100 mg seedlings using protein extraction buffer consisting of 50 mM Tris–HCl pH 7.5, 150 mMNaCl, 1% Triton X-100 and 1x Protease Inhibitor Cocktail (Sigma-Aldrich/Merck). Then the protein samples were mixed with SDS sample buffer and boiled for 5 min. The protein samples were separated on 10% SDS-polyacrylamide gels and separated proteins were transferred on to nitrocellulose membrane (BIORAD) for immunodetection. The membrane was immunoblotted using anti HA (Abcam, dilution 1:2000), anti MPK3, anti MPK4, anti MPK6 (Sigma-Aldrich, dilution 1:5000), anti-pTEpY (Cell signaling, dilution 1:2000) antibodies in TBST blocking buffer (50 mM Tris– HCl pH 8.0, 150 mM NaCl, 0.05% Tween-20, 5% dry skimmed milk) overnight. The immunoblotted membrane was washed with 1x TBST for three times for 10 min each and then incubated with anti-rabbit secondary antibody for 1.5 h. The membrane is again washed with TBST for three times. Chemiluminescent HRP Substrate (BIORAD) was used for visualization of secondary HRP bound antibodies.

### Bacterial protein expression and *in-vitro* phosphorylation assay

*MPK4* and *PIF4* were cloned in frame with 6xHis and GST tags, respectively. The proteins were induced in Rosetta strain of *E. coli* by addition of 1mM IPTG at 22℃. The recombinant MPK4 and PIF4 proteins with His and GST tags were purified by affinity chromatography using Ni-NTA and GST-sepharose beads, respectively. The total proteins were quantified by Bradford reagent (Sigma) and purity was analysed by SDS-PAGE. The *in-vitro* phosphorylation of PIF4 was performed as described previously (Raghuram et al., 2015).. MPK4 and PIF4 (1:10 enzyme substrate ratio) were incubated in the reaction buffer (20 mM Tris-HCl pH 7.5, 10 mM MgCl_2_, 25 μM ATP 1 μCi [γ-^32^P] ATP and 1 mM DTT) at 28℃ for 1 hr. The reaction was stopped using 6× SDS sample buffer, and the proteins were boiled for 5 min and loaded onto a 10% SDS-PAGE gel. After separation, gels were exposed to a phosphor screen and scanned using Typhoon FLA 9500 (GE Healthcare).

### *In Vivo* Phosphorylation Assay

*In Vivo* phosphorylation of PIF4 protein was examined by mobility sift assay using the Phos-tag reagent (NARD Institute) as described previously (Li et al., 2017).. Proteins were extracted from the seedlings grown at 22 and 28℃, and separated in 8% SDS-PAGE gel containing 50 μMPhos-tag and 100mM MnCl2. After electrophoresis, the gel was washed using the transfer buffer with 10 mM EDTA three times for 10 min each and finally with the transfer buffer without EDTA for 10 min, and then the separated proteins were transferred to nitrocellulose membrane. Anti HA antibody (Abcam) was used to detect HA tagged PIF4 protein.

### Construction of *pif4/mpk4* Double Mutants

*pif4/mpk4* double mutant lines were generated by crossing *pif4* mutant to *mpk4* heterozygous mutant plants as plants homozygous for the *mpk4* mutation produce very few buds. *mpk4* heterozygous mutant plants were used as female parent. 20 F1 seedlings with *PIF4/pif4//MPK4/mpk4* or *PIF4/pif4//MPK4/MPK4* genotypes were obtained by selection on basta containing 1/2MS medium since *pif4* mutant has T-DNA insertion with basta resistance gene. These 20 F1 plants were screened by PCR with WT *MPK4* & mutant *mpk4* allele specific primers to obtain heterozygous *PIF4/pif4//MPK4/mpk4* double mutant plants. Out of these 20, only 6 plants were found to be heterozygous *PIF4/pif4//MPK4/mpk4* double mutant (Supplemental Fig. S11A). These 6 F1 plants were self-pollinated to produce F2 population in which the two mutations were segregating. Homozygous *pif4/mpk4* double mutant F2 seedlings were easily identified due to their characteristic mutant phenotype when segregating F2 seeds were grown on MS medium. Further, these *pif4/mpk4* double mutant F2 plants were screened for the homozygosity of *pif4* and *mpk4* mutant alleles by PCR (Supplemental Fig. S11B). Primer sequences used in the screening of F1 and F2 plants have been mentioned in the Table S1. *pif4/mpk4* double mutant plants showed severely dwarf phenotype and did not produce viable seeds. Therefore, F2 plants which were homozygous mutant for *pif4* and heterozygous for *mpk4* (*pif4/pif4//MPK4/mpk4*), were screened (Supplemental Fig. S12) and self-pollinated to obtain homozygous double mutant seedlings for the analysis. Homozygous *pif4*/*mpk4* double mutant seedlings were identified on the basis of their distinct phenotype.

### Transient expression analysis in Arabidopsis seedlings

Transient expression in Arabidopsis seedlings was performed according to the method described previously with some variations (Verma & Burma, 2017).. The *Agrobacterium* cells carrying appropriate construct, were harvested by centrifugation of secondary culture at 6000 rpm for 15 min. *Agrobacterium* cell pellet was washed with washing solution (10 mM MgCl_2_,100 μMacetosyringone) two times. The *Agrobacterium* cells were again centrifuged and suspended into the co-cultivation medium (1 x Murashige-Skoog salts, 1 x B5 vitamins, 1% sucrose, 0.5g/L MES, 100 μM acetosyringone, 0.005% Silwet L-77, pH 6) to a final OD_600_ of 0.6.Then Arabidopsis seedlings were dipped this co-cultivation medium containing the *Agrobacterium* cells and incubated at 22℃ for 36 hrs. After incubation period, the seedlings were surface sterilized with surface sterilization solution (0.05% sodium hypochlorite) for 10 minutes followed by washing with water three times. Then, the seedlings were grown into 1/2 x MS medium at 22 & 28℃ for three days under SD conditions. Then, the seedlings were harvested and used for further analysis.

### Pi Staining and hypocotyl cells length measurement

Propidium Iodide (PI) powder (Sigma) was dissolved in PBS solution to prepare 5 mg/ml of stock solution. The stock solution was diluted to 5 μg/ml to prepare working solution. The seedlings were dipped into the solution for 10 min, then washed with water and then observed under the TSC SP8 confocal microscope (Leica). The excitation and image collection wavelengths were 405 nm and 410–503 nm, respectively. The Image J software was used for the measurement of cells length in the epidermis of middle portion of hypocotyl between shoot apical meristem and collet.

### Statistical analysis

The Student’s T-Tests and one-way ANOVA were performed to calculate significant differences which were indicated as ^∗^ P < 0.05, ^∗∗^ P < 0.005, and ^∗∗∗^ P < 0.0005. Details of statistical analysis and sample sizes in each experiment were mentioned in the Figure legends. ImageJ was used to quantify the intensities of bands in the western blot pics. Hypocotyl cells length were measured using ImageJ.

## Author contributions

NV and AKS planned the study and designed the experiments. NV, DS, LM, GB, and SN carried out the experiments. NV and AKS analyzed the data and wrote the manuscript. All the authors read and approved the final manuscript.

## Acknowledgement

NV is thankful to Department of Biotechnology (DBT) for fellowship under DBT-BioCare women scientist scheme. DS, LM, GB and SN are thankful to CSIR for fellowships. AKS thanks Sir JC Bose Fellowship from Science and Engineering Research Board, Department of Science and Technology, Government of India. The authors thank the Radioisotope facility and the Central Instrumentation Facility of NIPGR, New Delhi, India. The authors declare no conflict of interest.

## Supplemental data

**Supplemental Fig. S1.**
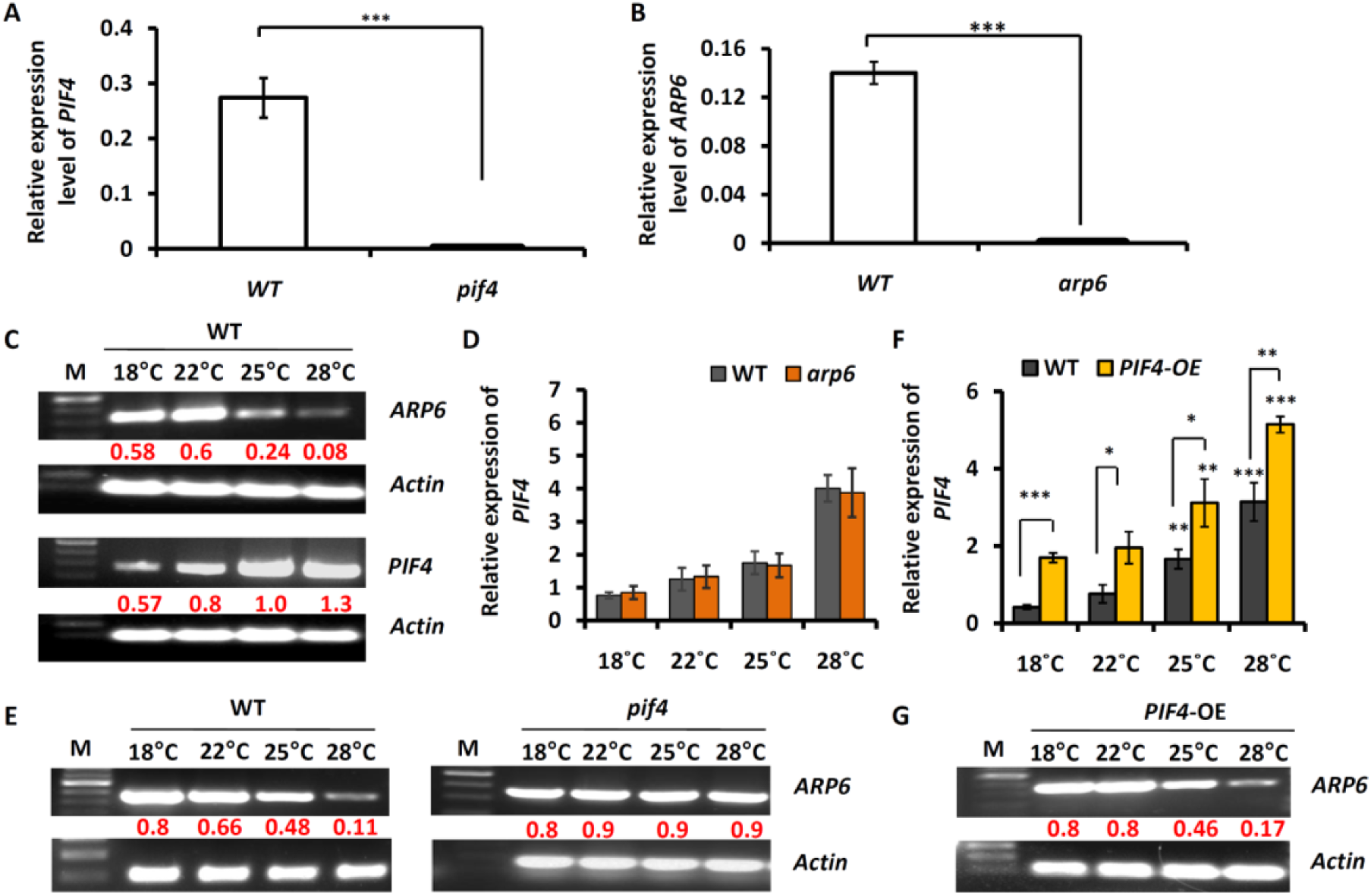
Analysis of genetic relationship between PIF4 and ARP6 (Supports Figure 1) **(A)** Bar graph showing the relative expression of *PIF4* in the seedlings of WT & *pif4* at 22℃. Each bar represents the mean ± SD of three independent experiments. Significant difference was obtained using Student’s t test; ***; *p* value < 0.0005. **(B)** Bar graph depicting the relative expression of *ARP6* in WT & *arp6* seedlings at 22℃. Each bar represents the mean ± SD of three independent experiments. Significant difference was calculated using Student’s t test; ***; *p* value < 0.0005. **(C)** Expression analysis of *ARP6* and *PIF4* in WT seedlings at 4 different temperatures by RT PCR (30 cycles for *ARP6* and 25 cycles for *PIF4*). Numbers in red colour depict the intensity of *ARP6* and *PIF4* transcript bands relative to *Actin*. **(D)** Bar graph showing the relative expression of *PIF4* in WT & *arp6* seedlings at 4 different temperatures. Each bar represents the mean ± SD of three independent experiments. The expression of *PIF4* at each temperature was compared between WT and *arp6*. No significant difference was obtained using Student’s t test; *p* value > 0.05. **(E)** Expression analysis of *ARP6* in WT and *pif4* seedlings at 4 temperatures using RT PCR. Numbers in red colour depict the intensity of *ARP6* transcript bands relative to *Actin*. **(F)** Bar graph showing the relative expression of *PIF4* in WT & *PIF4*-OE seedlings at 4 different temperatures. Each bar represents the mean ± SD of three independent experiments. The expression of *PIF4* in *PIF4*-OE seedlings was compared to its expression in WT at each temperature. Significant difference was calculated using Student’s t test. *; *p* value < 0.05, **; *p* value < 0.005, ***; *p* value < 0.0005. **(G)** Expression analysis of *ARP6* in *PIF4*-OE seedlings at 4 different temperatures by RT PCR. Numbers in red colour depict the intensity of *ARP6* transcript bands relative to *Actin*. 10 days old seedlings grown under SD conditions were used in each experiment.

**Supplemental Fig. S2.**
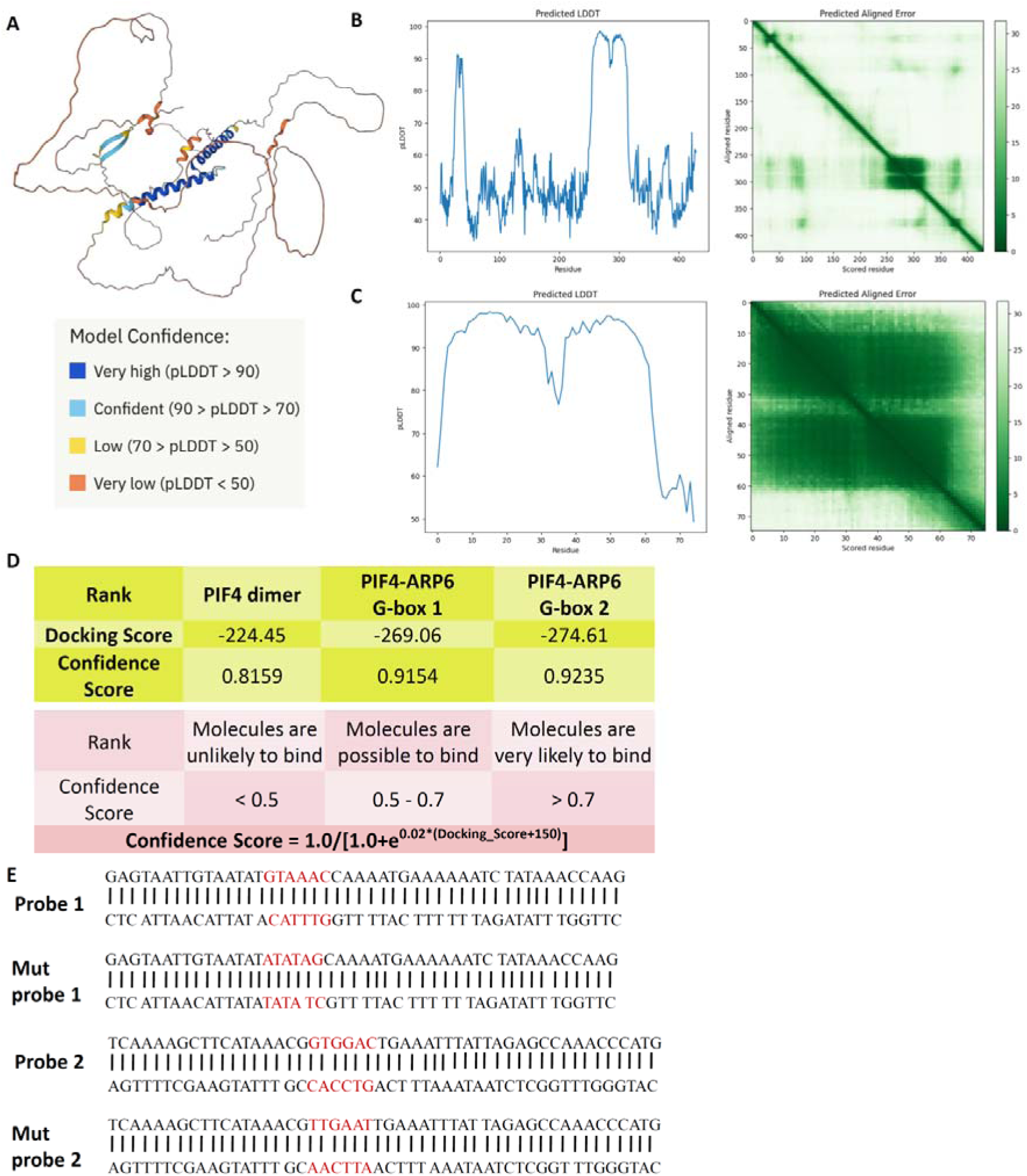
Prediction of PIF4 protein structure and sequences of probes used in EMSA (Supports Figure 2) **(A)** Predicted structure of PIF4 protein **(B)** Graphs showing the predicted LDDT and the predicted aligned error of the predicted structure of PIF4 protein. **(C)** Graphs showing the predicted LDDT and the predicted aligned error of the predicted structure of the bHLH domain of PIF4 protein. **(D)** Confidence score of the in-silico docking along with reference values. **(E)** Sequences of DNA probes used to perform EMSA.

**Supplemental Fig. S3.**
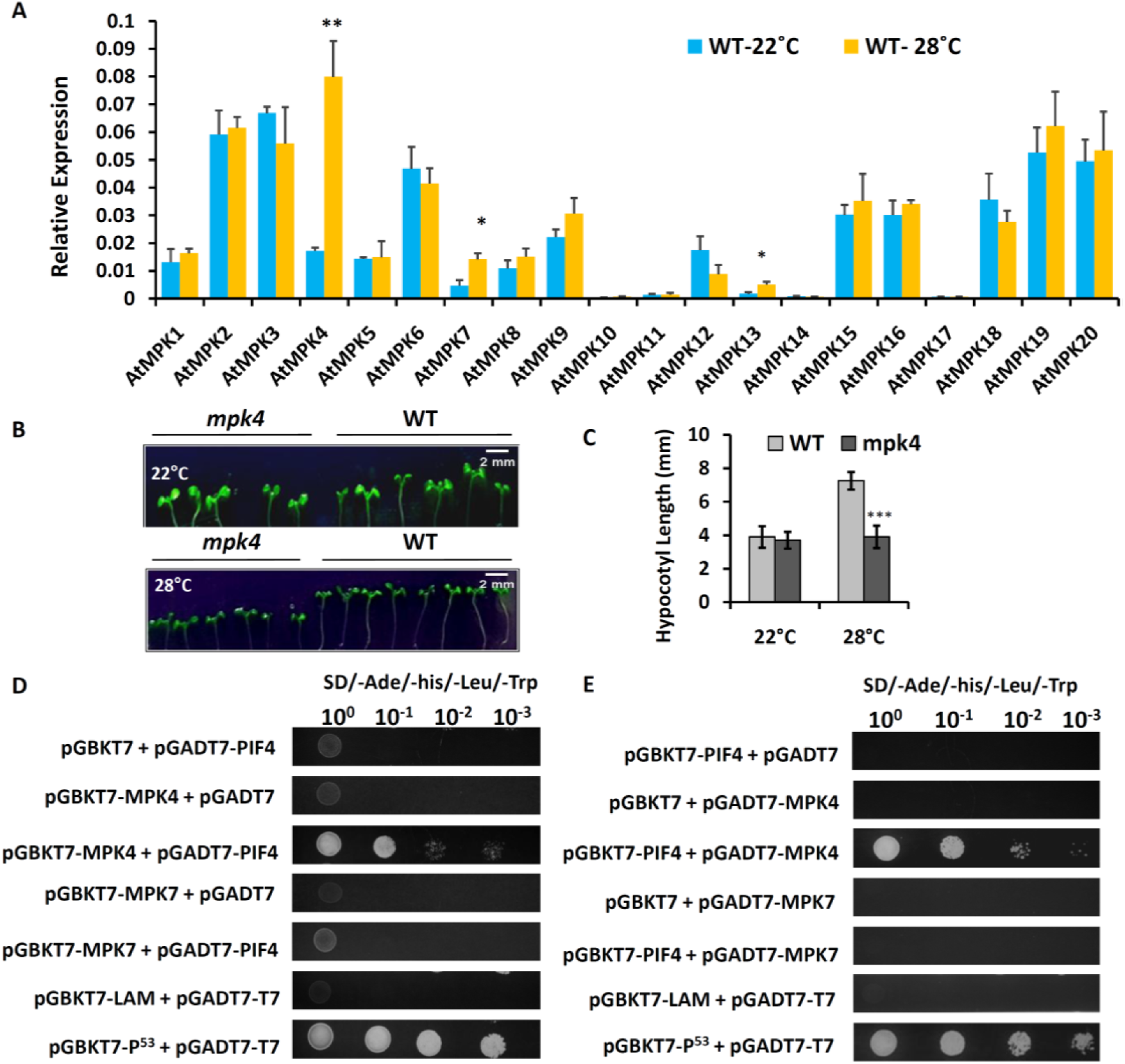
Upregulation of *MPK4* at elevated temperature and *in vitro* interaction of MPK4 *with PIF4* (Supports Figure 3) **(A)** Relative transcript level of MAP kinase genes (*MPK1*-*MPK20*) in WT Arabidopsis seedlings grown at 22 and 28℃. Each bar represents the mean ± s.d. of three independent replicates. Significant difference was calculated using Student’s t test. *; *p* value < 0.05, **; *p* value < 0.005 **(B)** Hypocotyl phenotype of 10 days old WT and *mpk4* seedlings grown under SD conditions at 22 and 28℃. **(C)** Quantitative measurement of hypocotyl length of WT and *mpk4* seedlings at 22 and 28℃. Error bars indicate s.d. (n = 20 plants). One-Way ANOVA: ***P<0.0005. **(D and E)** Y2H analysis with MAP kinases (MPK4 and MPK7) and PIF4 cloned into pGBKT7 and pGADT7 vectors (D), respectively, and pGADT7 and pGBKT7 vectors (E), respectively.

**Supplemental Fig. S4.**
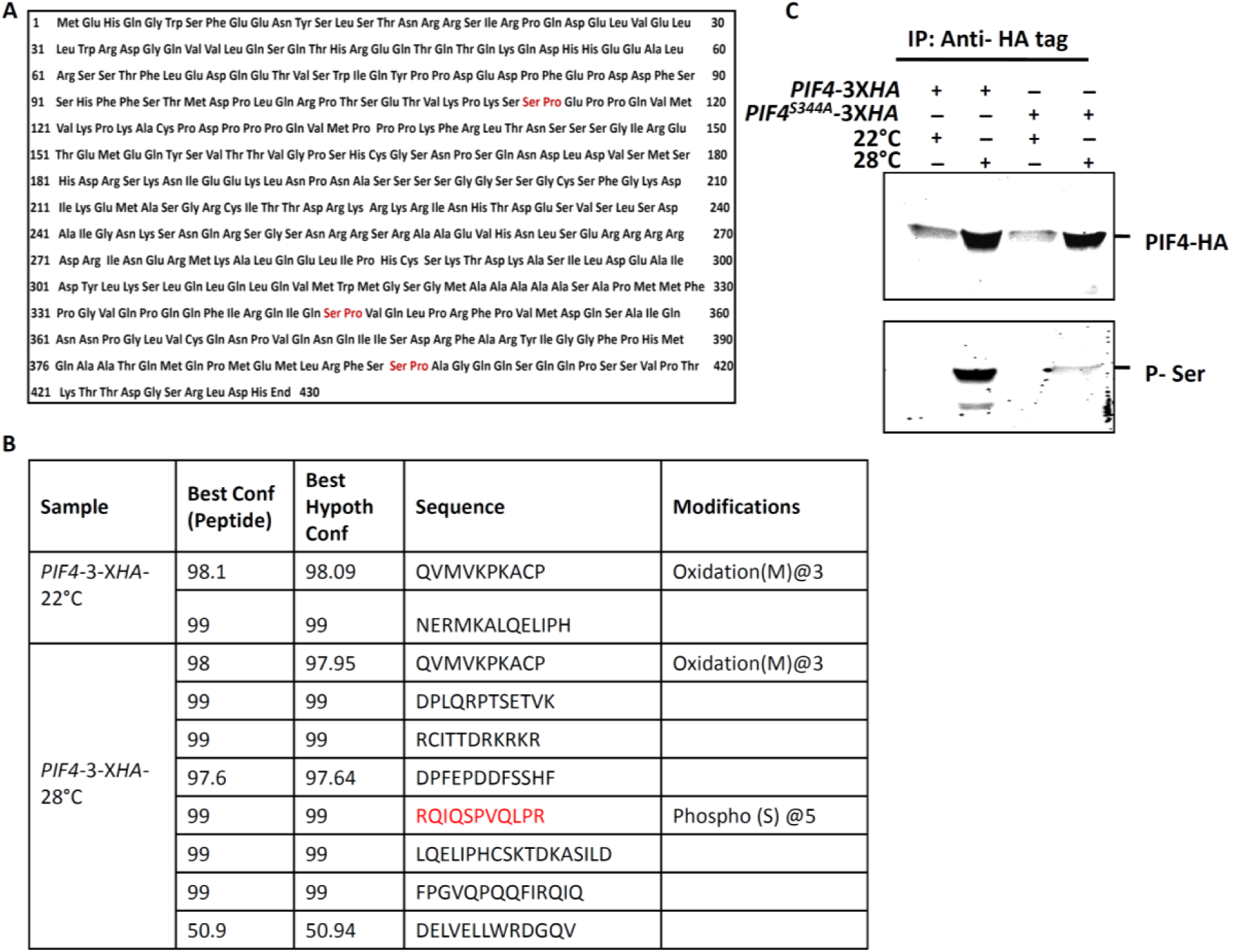
Phosphorylation of PIF4 at elevated temperature (Supports Figure 4) **(A)** Depiction of putative MAP kinase phosphorylation sites (shown in red color) consisting of serine residue followed by a characteristic proline amino acid in PIF4 protein sequence. **(B)** Distinct peptide summary of PIF4 protein immunoprecipitated from 10 days old p*PIF4-PIF4-*3X*HA* transgenic seedlings at 22 and 28℃ in IP-MS assays. Phosphopeptides (red colour) containing Ser344 residue followed by proline amino acid were detected in p*PIF4-PIF4-*3X*HA* transgenic seedlings grown at elevated temperature of 28℃. Best Conf (Peptide) determines the probability of the peptide fragments detected by the mass spectrum. **(C)** Co-IP assay showing the phosphorylation of WT PIF4 and phosphor-null PIF4^S344A^ proteins in a temperature-dependent manner using anti-phospho-serine (anti-pSer) antibody. *PIF4-*3X*HA* and *PIF4*^S344A^-3X*HA* transgenic seedlings grown at 22 and 28℃ were used for immunoprecipitation of both versions of PIF4 protein using anti-HA antibody. Precipitated proteins were analyzed by immunoblotting using anti-HA and anti-pSer antibodies, respectively.

**Supplemental Fig. S5.**
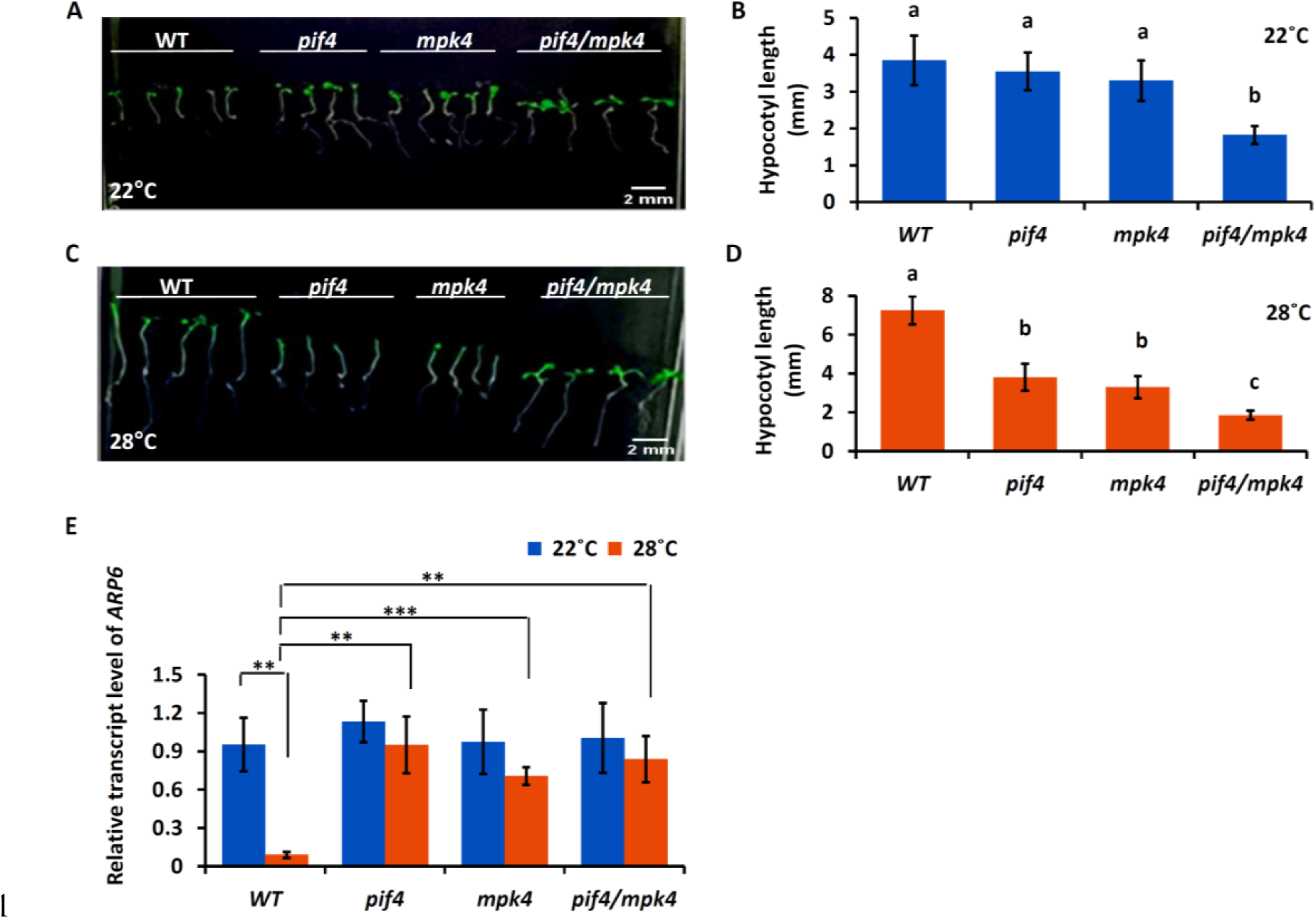
Analysis of hypocotyl length and *ARP6* expression in *pif4*/*mpk4* double mutant seedlings (Supports Figure 5) **(A** and **B)** Hypocotyl phenotype (A) and length (B) comparison of *pif4*/*mpk4* seedlings with WT, *pif4* & *mpk4* seedlings at 22 ℃. (B) Each bar represents the mean ± s.d. (n = 20). Letters above the bars indicate significant differences (P < 0.05), as calculated by One-way ANOVA with Tukey’s post hoc analysis. **(C** and **D)** Hypocotyl phenotype (C) and length (D) comparison of *pif4*/*mpk4* seedlings with WT, *pif4* & *mpk4* seedlings at 28 ℃. (D) Each bar represents the mean ± s.d. (n = 20). Letters above the bars indicate significant differences (P < 0.05), as calculated by One-way ANOVA with Tukey’s post hoc analysis. **(E)** Bar graph showing the relative transcript level of *ARP6* in 10 days old WT, *pif4*, *mpk4* & *pif4*/*mpk4* seedlings under SD conditions at 22 & 28℃. Each bar represents the mean ± s.d. of three independent experiments. The expression of *ARP6* at 28℃ was compared to its expression at 22℃ for each genotype and *ARP6* transcript levels in *pif4*, *mpk4* and *pif4*/*mpk4* were also compared to *ARP6* transcript level in WT at 22 and 28℃. Student’s t test. **; *p* value < 0.005, ***; *p* value < 0.0005.

**Supplemental Fig. S6.**
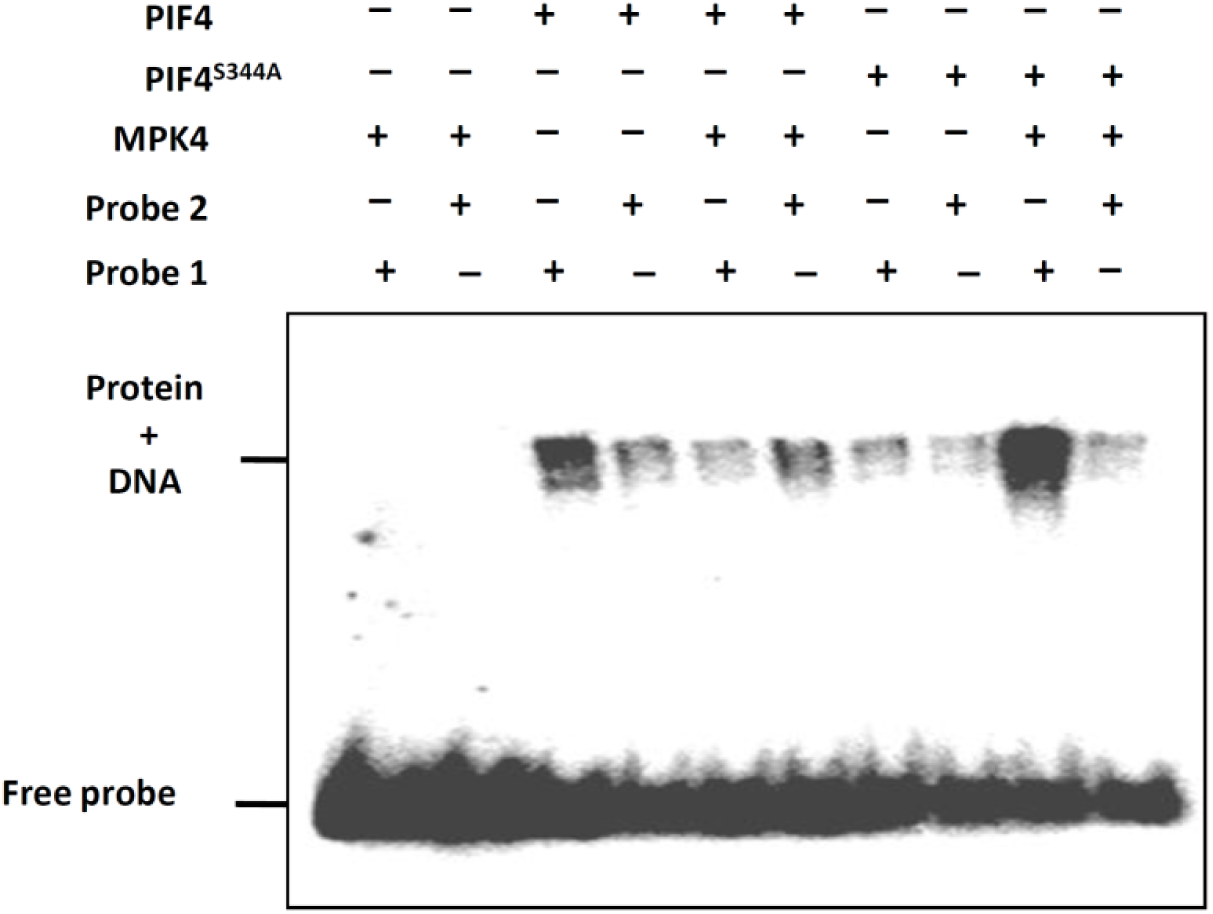
MPK4 mediated phosphorylation of PIF4 is not required for its binding to *ARP6* promoter (Supports Figure 5) EMSA showing the effect of MPK4 mediated phosphorylation on the interactions of PIF4 and *PIF4*^S344A^ proteins to *ARP6* promoter DNA probes.

**Supplemental Fig. S7.**
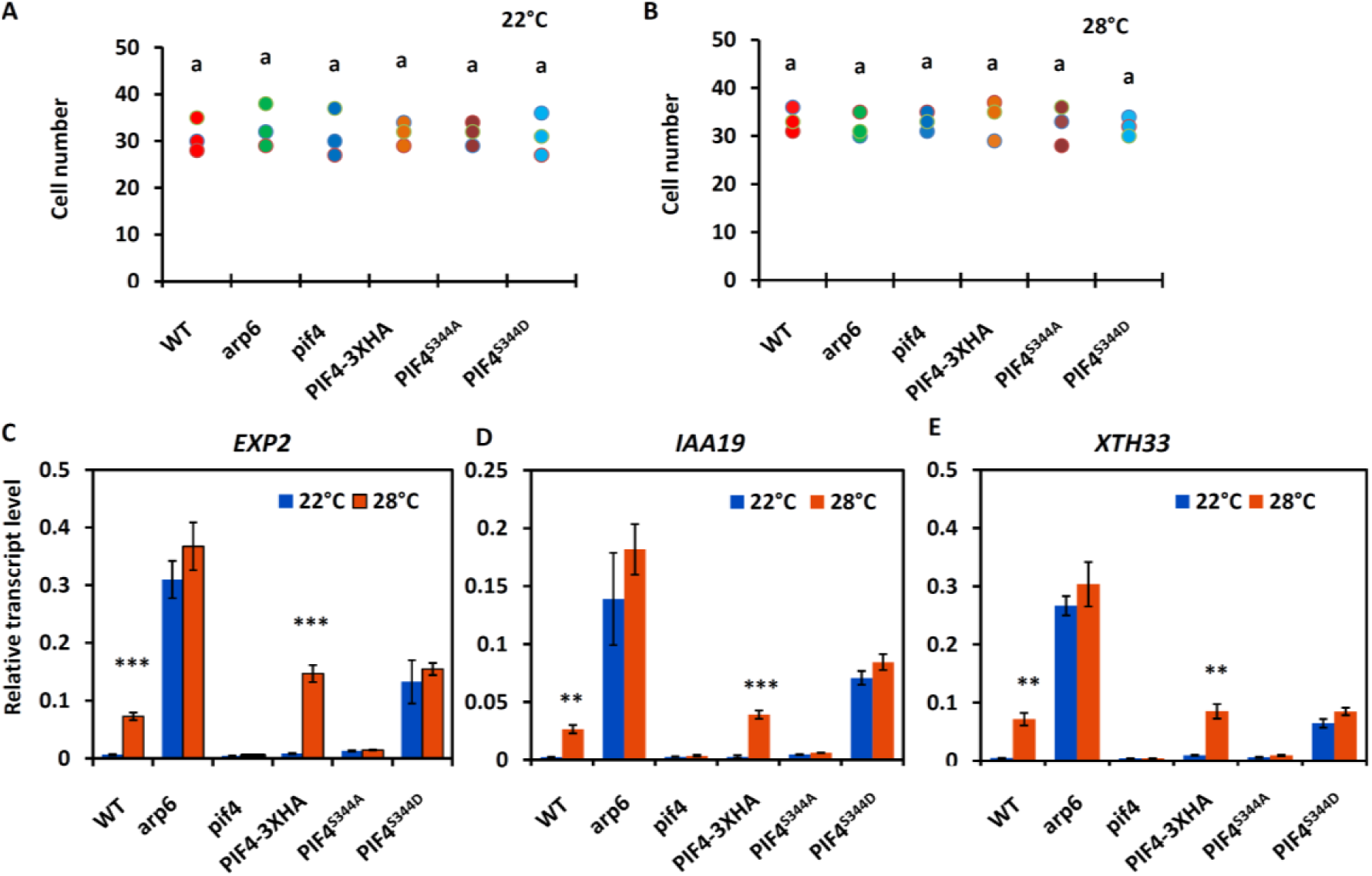
Effect of PIF4 phosphorylation on hypocotyl cell number and expression of genes involved in hypocotyl elongation at 22 and 28℃ (Supports Figure 7) **(A)** Total cells number in the hypocotyls of 10 days old WT, *arp6*, *pif4*, *PIF4-*3X*HA*, *PIF4*^S344A^ and *PIF4*^S344D^ seedlings at 22℃. **(B)** Total cells number in the hypocotyls of 10 days old WT, *arp6*, *pif4*, *PIF4-*3X*HA*, *PIF4*^S344A^ and *PIF4*^S344D^ seedlings at 28℃. n = 3 hypocotyls **(C, D and E)** Quantification of *EXP2* (C), *IAA19* (D) and *XTH33* (E) transcript levels in the 10 days old WT, *arp6*, *pif4*, *PIF4-*3X*HA*, *PIF4*^S344A^ and *PIF4*^S344D^ seedlings at 22℃ and 28℃. Each bar represents the mean ± s.d. of three independent experiments. The expression of all the three genes at 28℃ was compared to its expression at 22℃ for each genotype. Student’s t test was used to calculate the significant difference. (**; *p* value < 0.001, ***; *p* value < 0.0001)

**Supplemental Fig. S8.**
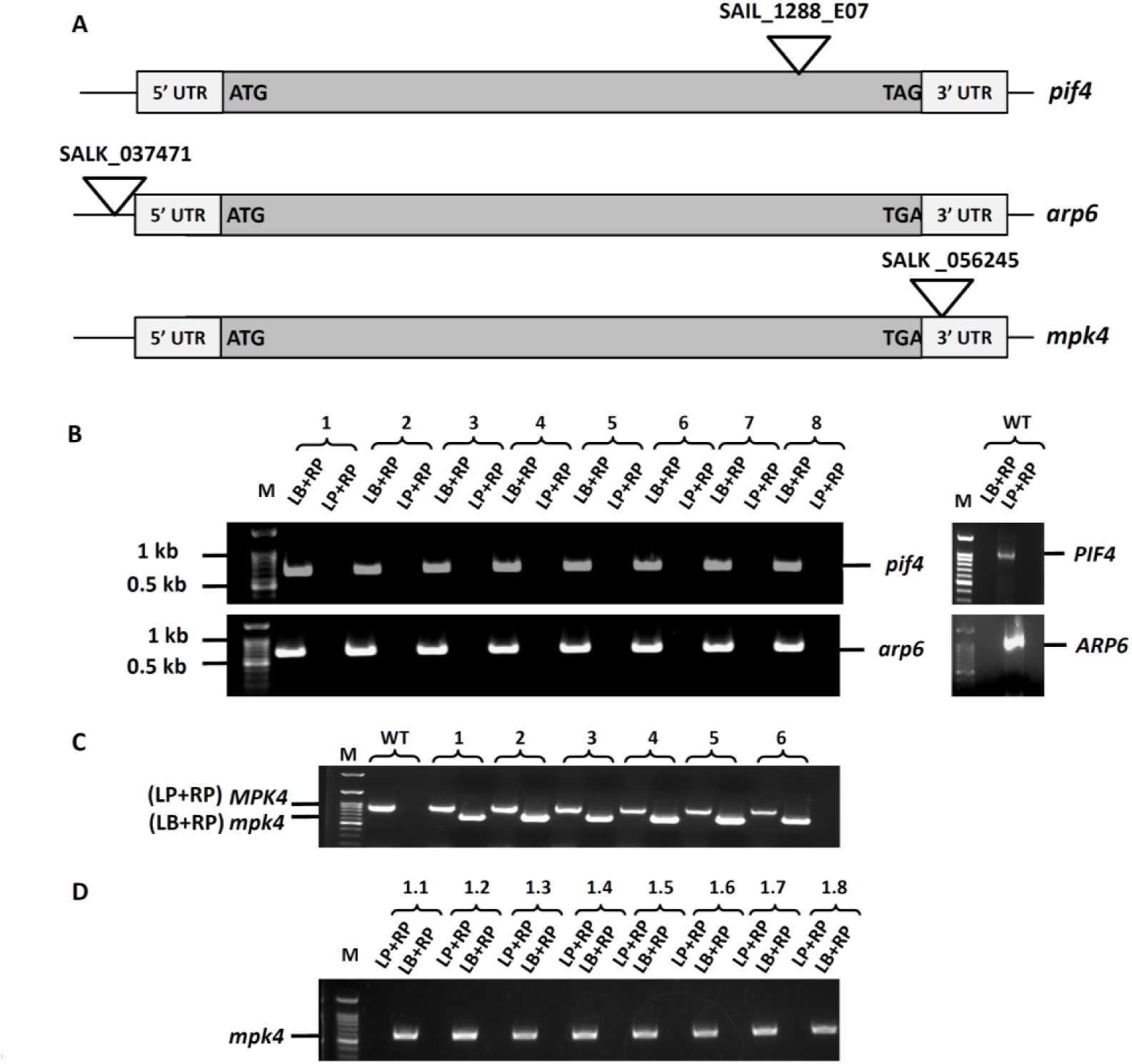
Screening of *pif4*, *arp6 and mpk4* plants by PCR genotyping (Supports Figure 1, 3, 4, 5, 6, 7 and 8) **(A)** Schematic diagrams of T-DNA insertion sites in *pif4, arp6 and mpk4* mutants. The 5’ UTR, start codon (ATG), the stop codon (TAG/ TGA) and 3’ UTR are indicated. **(B)** PCR genotyping of 8 plants of *pif4* & *arp6* with mutant allele (LB+RP) and WT allele (LP+RP) specific primers. Both the genotypes, *pif4* & *arp6* showed homozygosity of *pif4* and *arp6* alleles, respectively. **(C & D)** Screening of *mpk4* plants by PCR genotyping (C) PCR genotyping of 6 heterozygous MPK4/*mpk4* plants grown from the seeds procured from ABRC with mutant allele (LB+RP) and WT allele (LP+RP) specific primers. All the tested plants were found to be heterozygous for *mpk4* mutation. (D) Screening of 8 plants with distinct phenotype, grown from the seeds produced by the self pollination of heterozygous *mpk4* plant-1, using mutant allele (LB+RP) and WT allele (LP+RP) specific primers. All the tested plants showed homozygosity of *mpk4* allele. I00 bp ladder was used as a size marker in all PCR genotyping experiments.

**Supplemental Fig. S9.**
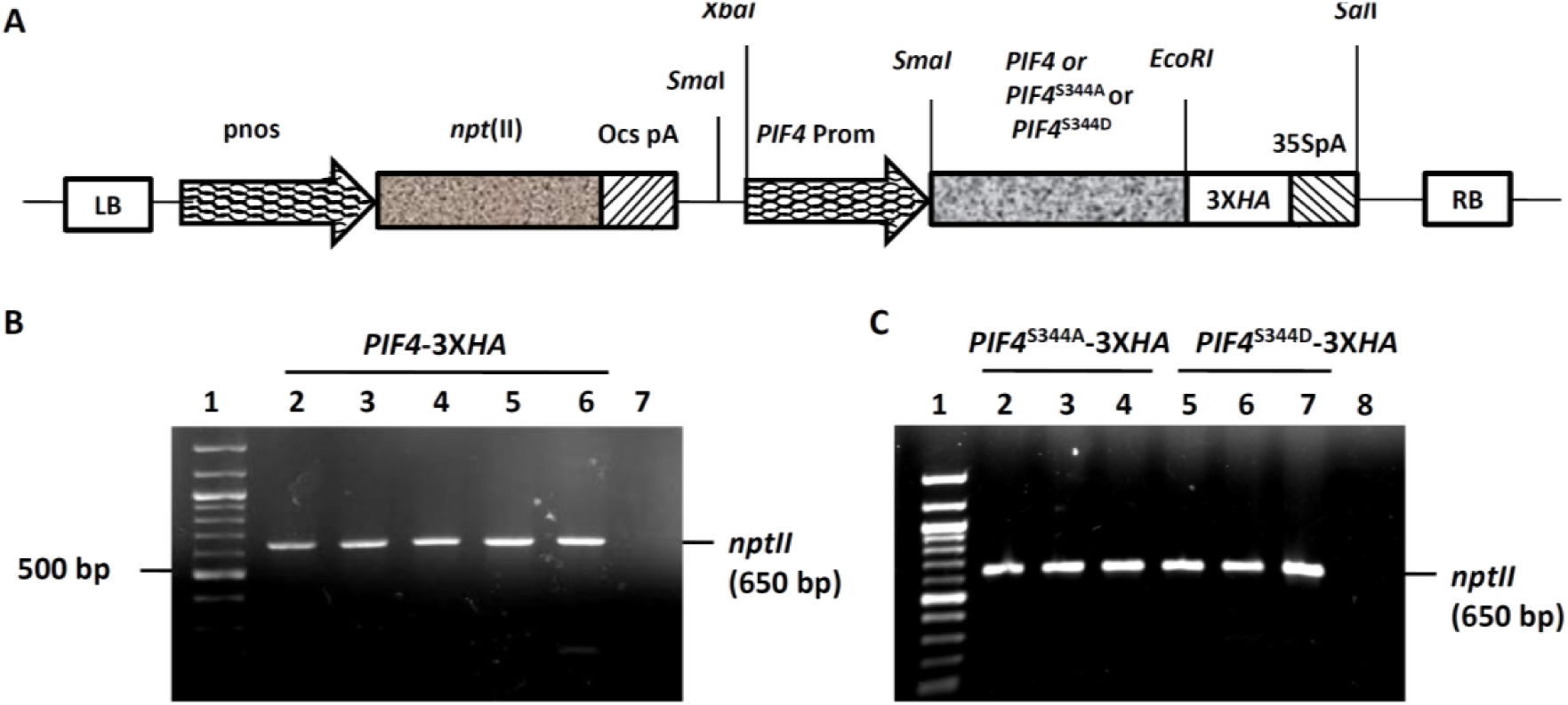
Development of *PIF4*-3X*HA*, *PIF4*^S344A^-3X*HA* and *PIF4*^S344AD^-3X*HA* transgenic plants. (Supports Figure 2, 3, 4, 5, 6, and 7) **(A)** Schematic showing the construct design of WT, phosphor-null (*PIF4*^S344A^) and phosphor-mimetic (*PIF4*^S344D^) versions of *PIF4* between the LB and RB of T-DNA in the binary vector. WT or mutated *PIF4* gene cDNA with 3 copies of *HA tag* were fused downstream to the *PIF4* promoter. Selection marker expression cassette coding for kanamycin resistance, carries *nos* promoter driven *nptII* gene. **(B & C)** Confirmation of *PIF4*-3X*HA*, *PIF4*^S344A^-3X*HA* and *PIF4*^S344AD^-3X*HA* transgenic lines by PCR amplification. (B) Gel picture showing showing PCR amplification of *nptII* gene of kanamycin resistant *PIF4*-3X*HA* transgenic F1 plants. Lane 1; I00 bp ladder was used as size marker. lanes 2, 3, 4, 5 & 6; Amplification of *nptII* gene & Lane 7; Negative control (amplification using genomic DNA of *pif4* mutant plant). (C) Gel picture showing PCR amplification of *nptII* gene of kanamycin resistant, *PIF4*^S344A^-3X*HA* and *PIF4*^S344AD^-3X*HA* transgenic F1 plants. Lane 1; I00 bp ladder was used as size marker. lanes 2, 3, & 4; Amplification of *nptII* gene from *PIF4*^S344A^-3X*HA* plants, lanes 5, 6, & 7; Amplification of *nptII* gene from *PIF4*^S344D^-3X*HA* plants & Lane 8; Negative control (amplification using genomic DNA of *pif4* mutant plant).

**Supplemental Fig. S10.**
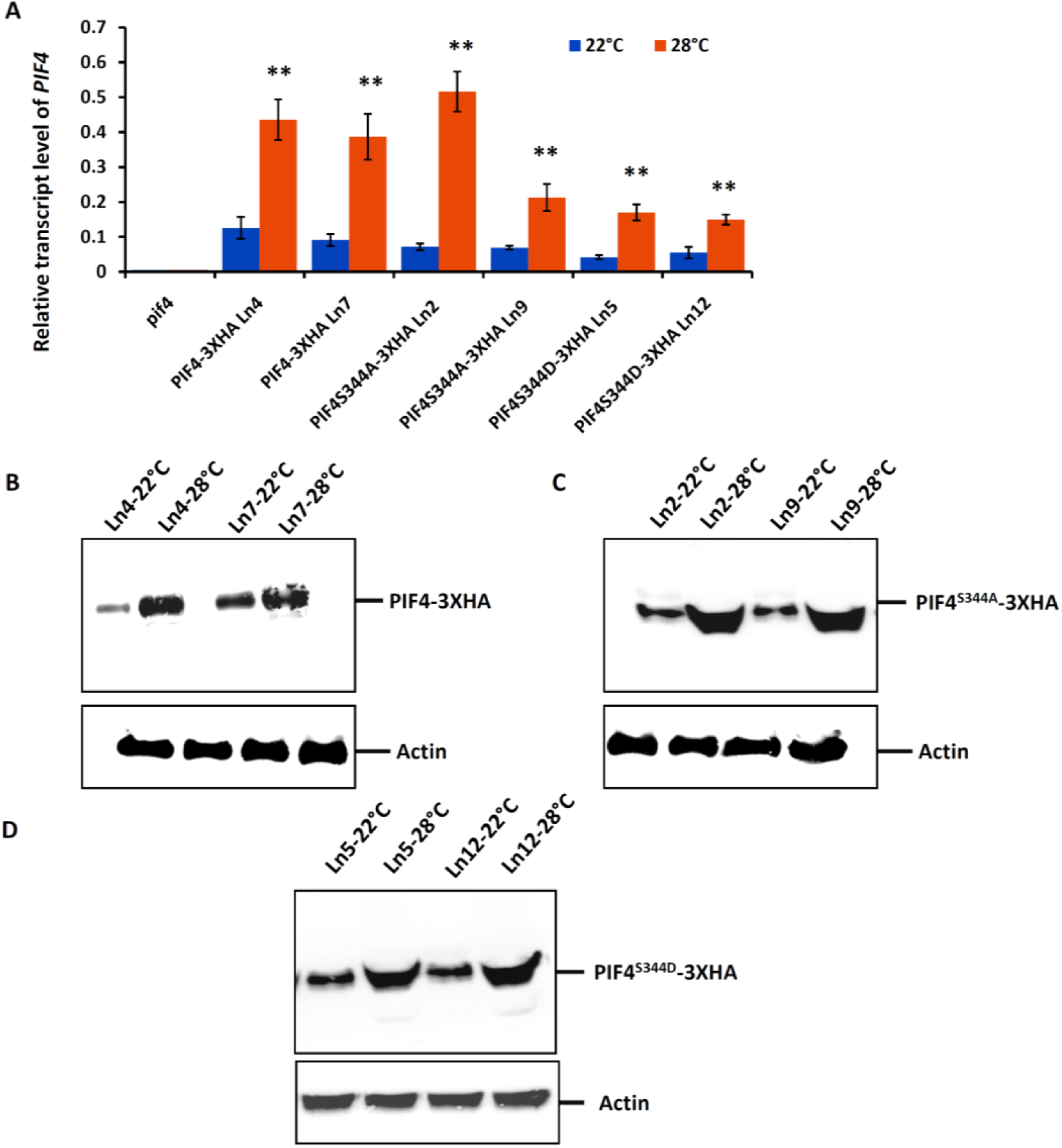
Characterization of WT *PIF4*-3X*HA*, phosphor-null *PIF4*^S344A^-3X*HA* and phosphor-mimetic *PIF4*^S344D^-3X*HA* transgenic lines. (Supports Figure 2, 3, 4, 5, 6, and 7) **(A)** Bar graph showing the transcript level of *PIF4*, *PIF4*^S344A^ and *PIF4*^S344D^ in 10 days old *pif4* and WT or various phosphosite mutated homozygous *PIF4* complementation lines at 22 and 28℃. Error bars represent SD (n = 3). **(B)** Protein level of PIF4 in the seedlings of homozygous *PIF4*-3X*HA* complementation lines at 22 and 28℃. **(C)** Protein level of PIF4^S344A^ in the seedlings of homozygous *PIF4*^S344A^ -3×*HA* complementation lines at 22 and 28℃. **(D)** Protein level of PIF4^S344D^in the seedlings of homozygous *PIF4*^S344D^-3X*HA* complementation lines at 22 and 28℃. Immunoblotting using anti-Actin antibody was used as loading control.

**Supplemental Fig. S11.**
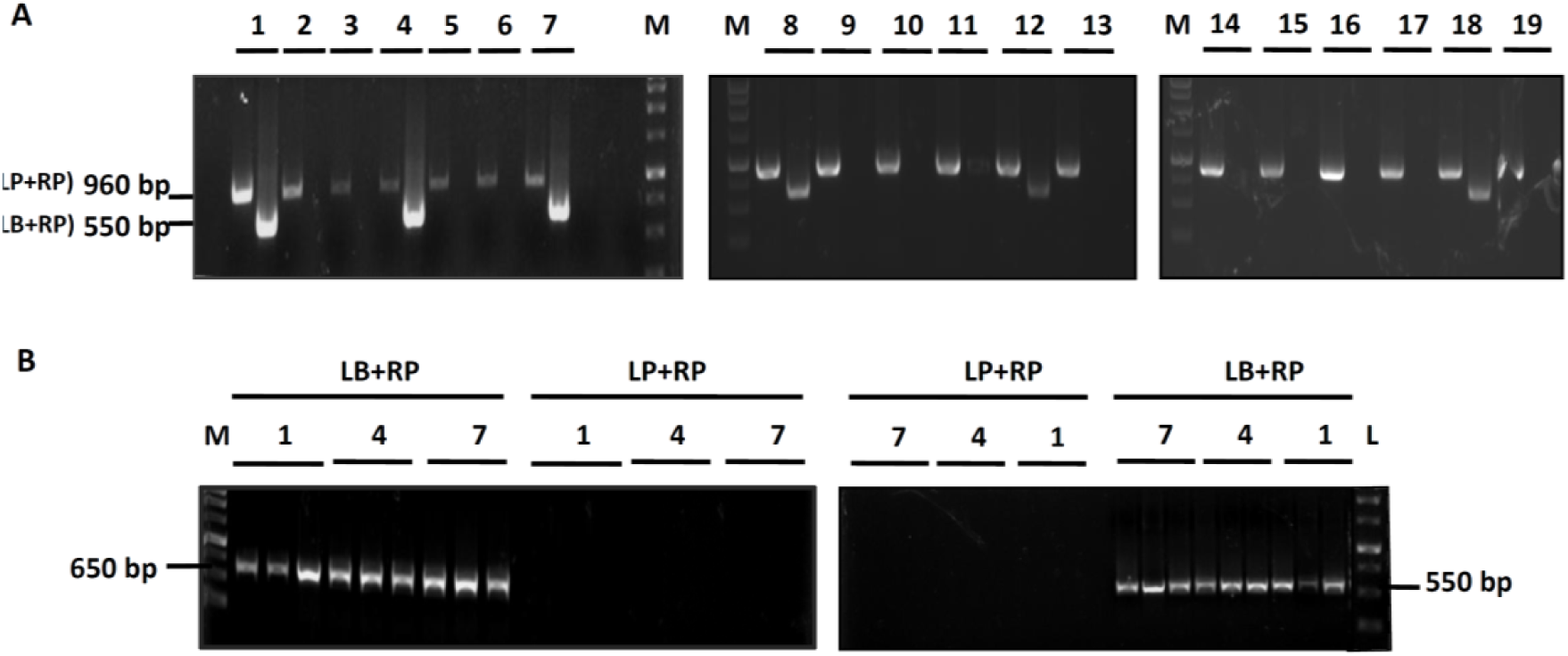
Screening of *pif4*/*mpk4* double mutant by PCR genotyping. (Supports Figure 5) **(A)** PCR genotyping of *pif4*/*mpk4* heterozygous F1 plants. PCR screening of 20 basta resistant F1 plants with *MPK4* (LP+RP) and *mpk4* (LB+RP) alleles specific primers. Out of 20, lines 1, 4, 7, 8, 12 & 18 were screened to be *pif4*/*mpk4* heterozygous double mutant. I kb ladder was used as a size marker. **(B)** PCR genotyping of *pif4*/*mpk4* homozygous F2 plants. PCR screening of F2 seedlings with distinct phenotype to confirm the homozygosity of *pif4* and *mpk4* mutant alleles. LB+RP for *pif4* and *mpk4* alleles produced PCR products of 650 and 550 bp, respectively. I kb ladder was used as a size marker.

**Supplemental Fig. S12.**
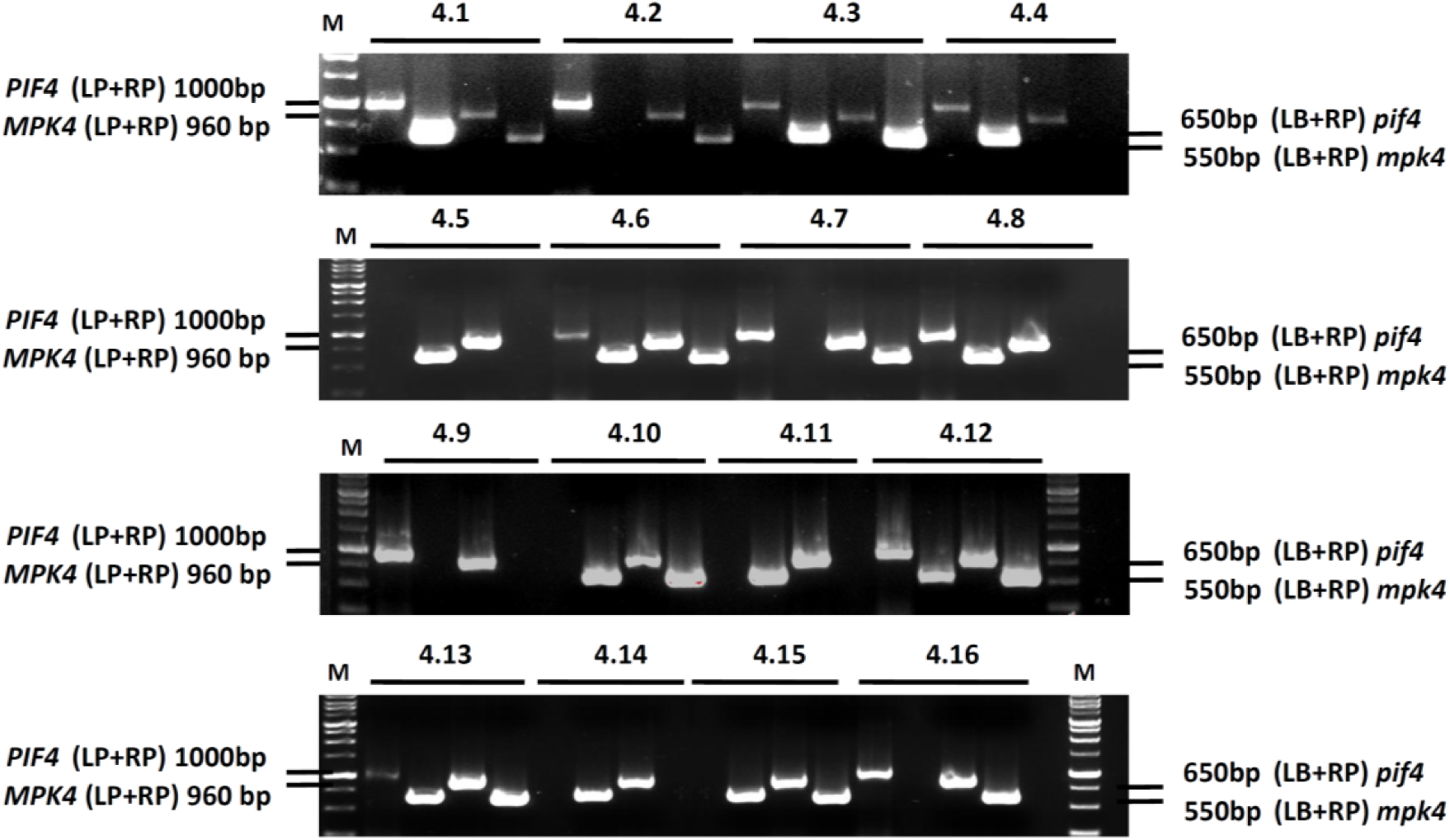
Screening of *pif4/pif4//MPK4/mpk4* F2 plants by PCR genotyping. (Supports Figure 5) PCR genotyping of F2 seedlings to screen for the plants homozygous for *pif4* mutation and heterozygous for *mpk4* mutation. LB+RP for *pif4* and *mpk4* alleles produced PCR products of 650 and 550 bp, respectively. LP+RP for *PIF4* and *MPK4* alleles produced PCR products of 960 and 1000 bp, respectively. I kb ladder was used as size marker. F2 plants, 4.10 & 4.15 are homozygous for *pif4* and heterozygous for *mpk4* allele.

## Supplemental tables

**Supplemental Table S1.**
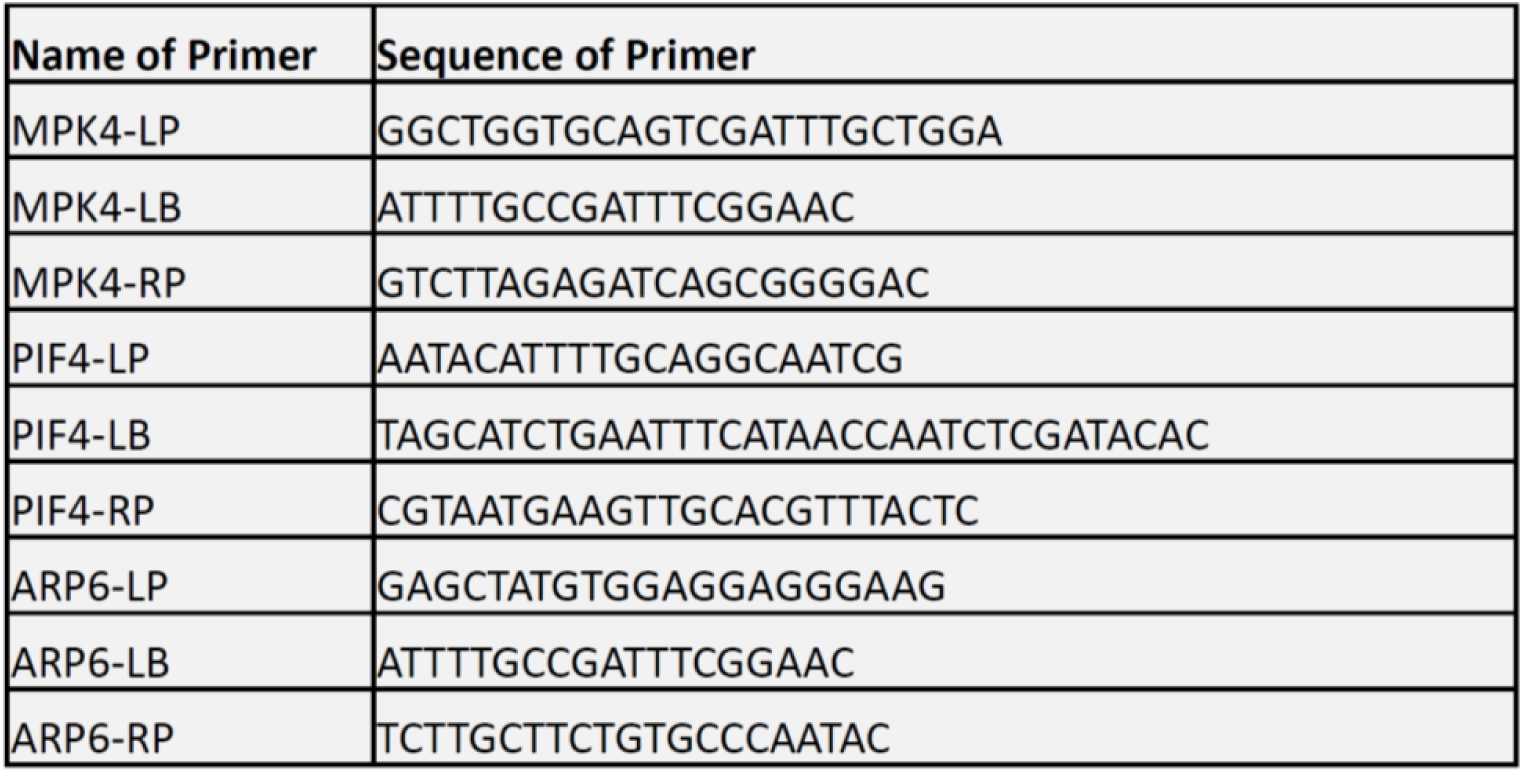
Sequences of primers used to screen the Arabidopsis mutants

**Supplemental Table S2.**
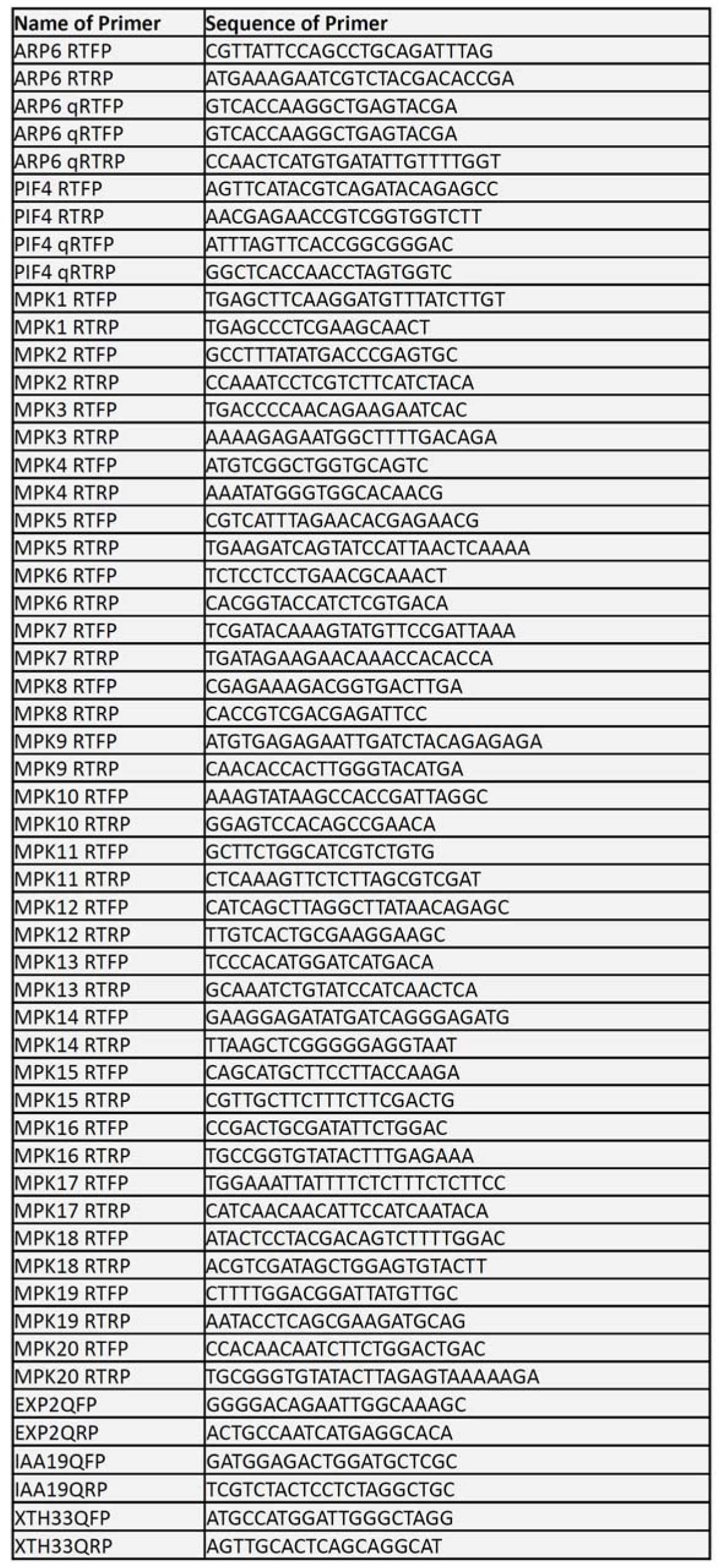
Sequences of primers used in the genes expression analysis

**Supplemental Table S3.**
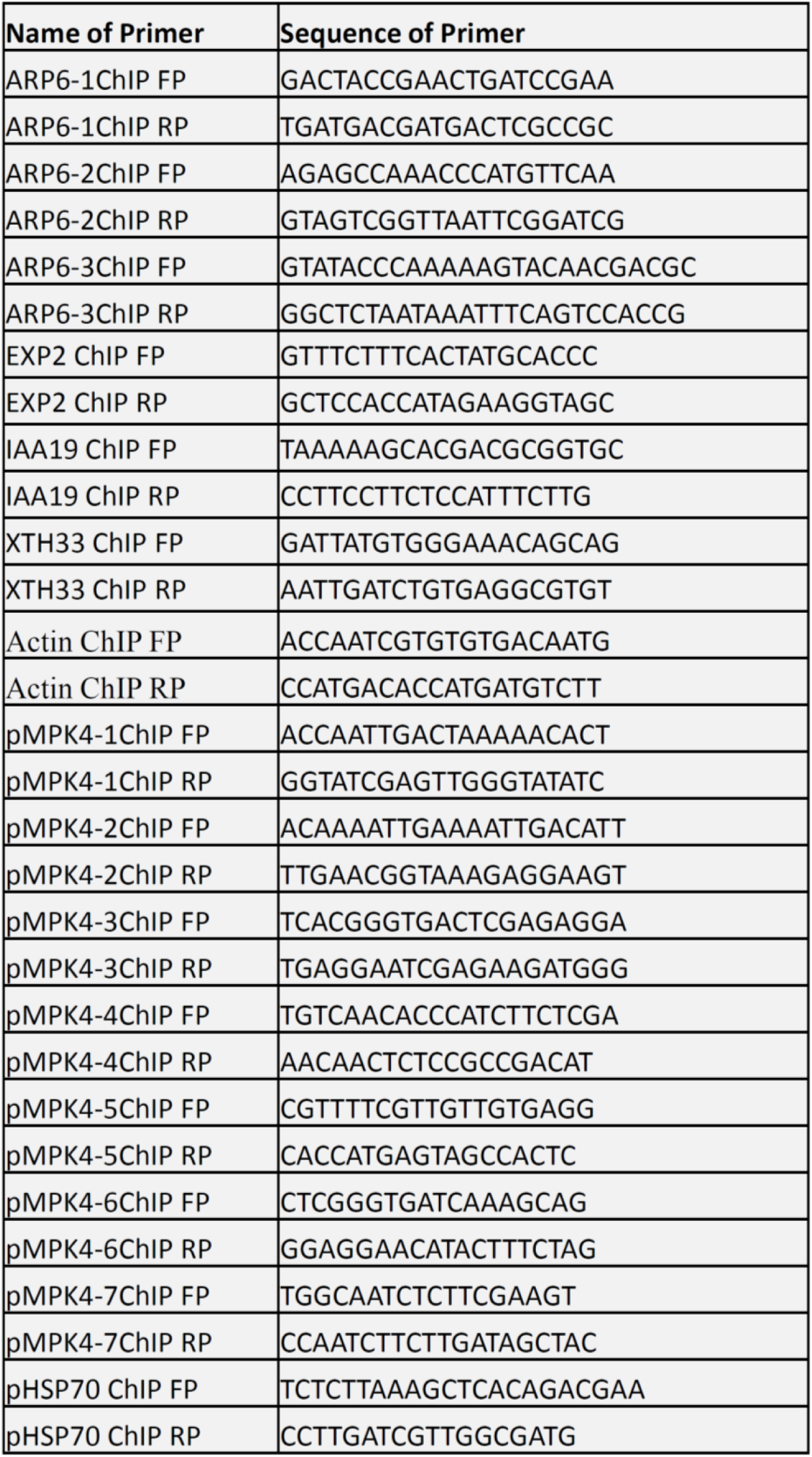
Sequences of primers used in ChIP-qPCR

